# Emergence of the Ug99 lineage of the wheat stem rust pathogen through somatic hybridisation

**DOI:** 10.1101/692640

**Authors:** Feng Li, Narayana M. Upadhyaya, Jana Sperschneider, Oadi Matny, Hoa Nguyen-Phuc, Rohit Mago, Castle Raley, Marisa E. Miller, Kevin A.T. Silverstein, Eva Henningsen, Cory D. Hirsch, Botma Visser, Zacharias A. Pretorius, Brian J. Steffenson, Benjamin Schwessinger, Peter N. Dodds, Melania Figueroa

**Author notes:** These authors contributed equally to this work (Feng Li, Narayana M. Upadhyaya). Correspondence to: Melania Figueroa, Peter N. Dodds.

## Abstract

Parasexuality contributes to diversity and adaptive evolution of haploid (monokaryotic) fungi. However non-sexual genetic exchange mechanisms are not defined in dikaryotic fungi (containing two distinct haploid nuclei). Newly emerged strains of the wheat stem rust pathogen, *Puccinia graminis* f. sp. *tritici* (*Pgt*), such as Ug99, are a major threat to global food security. Here we show that Ug99 arose by somatic hybridisation and nuclear exchange between dikaryons. Fully haplotype-resolved genome assembly and DNA proximity analysis revealed that Ug99 shares one haploid nucleus genotype with a much older African lineage of *Pgt*, with no recombination or reassortment. Generation of genetic variation by nuclear exchange may favour the evolution of dikaryotism by providing an advantage over diploidy.

---

Generation of genetic diversity is crucial for the evolution of new traits, with mutation and sexual recombination as the main drivers of diversity in most eukaryotes. However, many species in the fungal kingdom can propagate asexually for extended periods and therefore understanding alternative mechanisms contributing to genetic diversity in asexual populations has been of great interest^1,2^. Some fungi can use a parasexual mechanism to exchange genetic material independently of meiosis^2^. This process involves anastomosis of haploid hyphae and fusion of two nuclei to generate a single diploid nucleus, which subsequently undergoes progressive chromosome loss to generate recombinant haploid offspring. Parasexuality has been described in members of the ascomycete phylum (64% of described fungal species) in which the dominant asexually propagating form is haploid^3^. However, in basidiomycete fungi (34% of described species), the predominant life stage is generally dikaryotic, with two different haploid nuclei maintained within each individual^3^. The role of non-sexual genetic exchange between such dikaryons in generating genetic diversity is not known.

Basidiomycetes include many fungi with critical ecosystem functions, such as wood decay and plant symbiosis, as well as agents of important human and plant diseases^1^. Rust fungi (subphylum Pucciniomycotina) comprise over 8,000 species including many pathogens of major agricultural and ecological significance^4^. These organisms are obligate parasites with complex life cycles that can include indefinite asexual reproduction through infectious dikaryotic urediniospores. Early researchers speculated that rust fungi can exchange genetic material during the asexual phase^5-8^, but these hypotheses could not be confirmed molecularly. Some naturally occurring rust pathotypes have been suggested to have arisen by somatic hybridisation and genetic exchange based on limited molecular evidence of shared isozyme or random amplified polymorphic DNA (RAPD) markers^9,10^. Mechanisms underlying genetic exchange are unknown, but may involve hyphal anastomosis followed by nuclear exchange and/or nuclear fusion and recombination^11^. Recent advances in assembling complete karyon sequences in rust fungi^12,13^ provide the opportunity to definitively detect and discriminate between nuclear exchange and recombination.

The Ug99 strain (race TTKSK) of the wheat stem rust pathogen *Puccinia graminis* f. sp. *tritici* (*Pgt*) presents a significant threat to global wheat production^14^. It was first detected in Uganda in 1998 and described in 1999^15^, and has since given rise to an asexual lineage that has spread through Africa and the Middle East causing devastating epidemics^14^. The origin of the Ug99 lineage is unknown, although it is genetically distinct from other *Pgt* races^16,17^. To resolve the genetic makeup of Ug99, we generated a haplotype-phased genome reference for the original Ug99 isolate collected in Uganda^15^. In addition, we also generated a similar reference for an Australian *Pgt* isolate of pathotype 21-0^18,19^. This isolate is a member of a longstanding asexual lineage that has been predominant in southern Africa since the 1920’s and spread to Australia in the 1950’s^19-21^.

## Results

### Haplotype phased genome assembly

We generated polished long-read genome assemblies for both Ug99 (Supplementary Table 1) and *Pgt*21-0 using single-molecule real-time (SMRT) and Illumina sequence data (Supplementary Tables 2 and 3). The assemblies (177 and 176 Mbp, respectively, Supplementary Table 4) were twice the size of a collapsed haploid assembly for a North American *Pgt* isolate^22^. This suggested that the sequences of the two haploid karyons in each isolate were represented independently. Both genomes contained over 96% of conserved fungal genes, and the *Pgt*21-0 assembly contained 69 telomeres (Supplementary Table 4), out of an expected 72 in a dikaryotic genome with n=1^23^. We developed a gene synteny approach to identify sequences representing alternate haplotypes within each assembly (Fig. 1), which were assigned to bins containing homologous pairs of sequences from each haplotype. The 44 bins in *Pgt*21-0 and 62 bins in Ug99 represented 95% and 94% of the respective assemblies (Supplementary Tables 4 and 5).

**Fig. 1.**
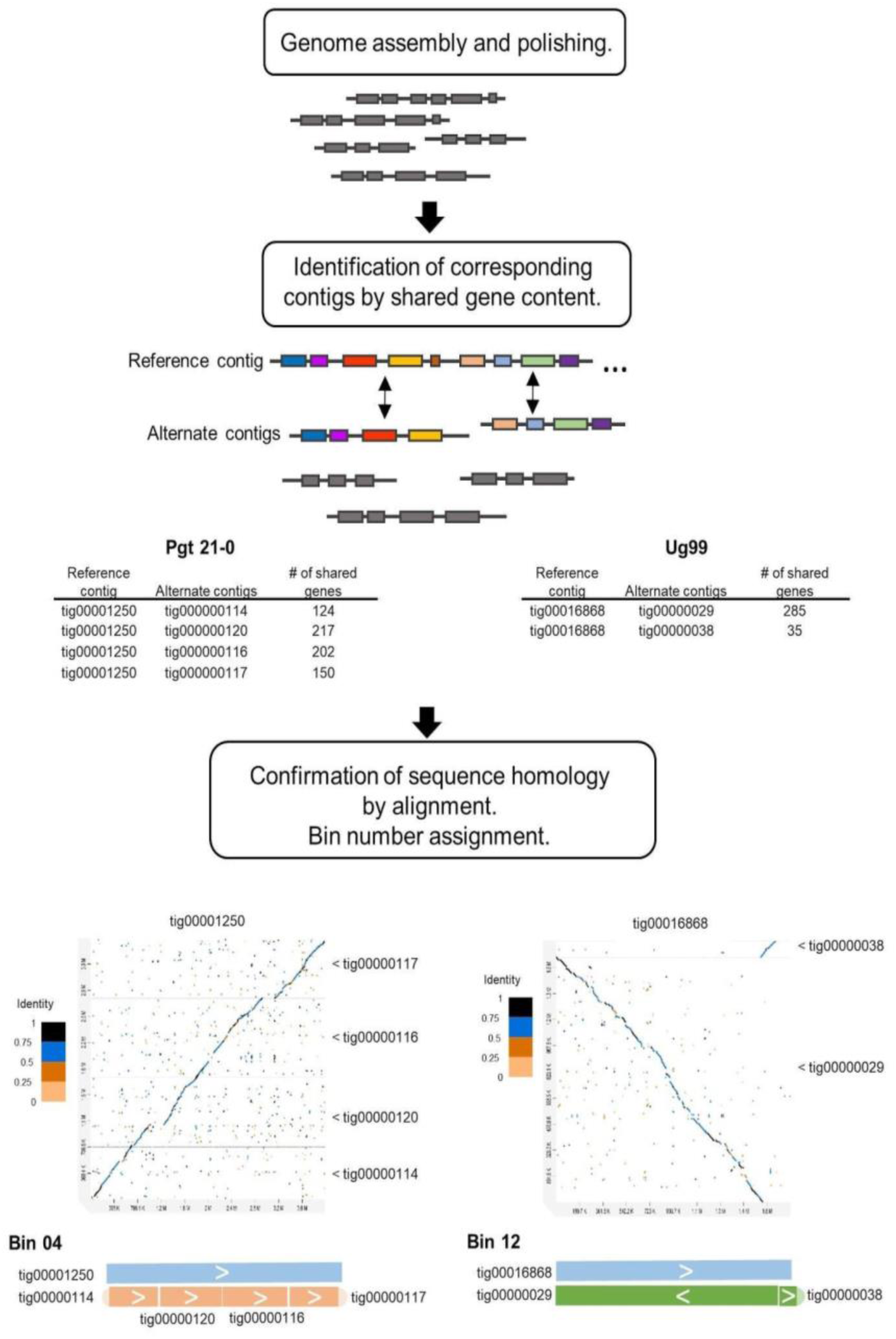
Strategy to identify homologous contigs and de-duplicate genome assemblies based on gene synteny. Shared gene content between contigs was assessed by alignment of 22,484 predicted genes to the full genome assemblies and the contig positions of the top two hits of each gene were recorded (represented as rectangle boxes on the contigs). Contigs containing at least five shared genes were considered as potential haplotype pairs. Sequence collinearity between such putative alternate contigs was assessed by alignment, and homologous matching contigs were assigned to Bins. Examples shown are for Bin 04 and Bin 12 from *Pgt*21-0 and Ug99 respectively.

The *AvrSr50* and *AvrSr35* genes encode dominant avirulence factors recognized by wheat resistance genes^24,25^. These two genes are located in close proximity to each other and both haplotypes of this locus were assembled as alternate contigs in *Pgt*21-0 and Ug99 (Fig. 2a). Both isolates were heterozygous for *AvrSr50* with one allele containing a ∼ 26 kbp-insertion. *Pgt Pgt*21-0 was also heterozygous for *AvrSr35*, with one allele containing a 400 bp MITE insertion previously described^25^. Although PCR amplification had identified only a single *AvrSr35* allele in Ug99 suggesting homozygosity^25^, a second allele identified in the Ug99 genome assembly contained a 57 kbp insertion that would have prevented its PCR amplification. The presence of the insertion was supported by read alignments across this region and confirmed by DNA amplification and amplicon sequencing of flanking border regions (Supplementary Fig. 1). Thus, Ug99 is also heterozygous for avirulence on *Sr35*, and may therefore mutate to virulence on this wheat resistance gene more readily than if it were homozygous. This is an important finding as it will inform *Sr35* deployment strategies against Ug99. Strikingly, the *AvrSr35/virSr50* haplotype of this locus is very similar (>99% sequence identity) in Ug99 and *Pgt*21-0, while the two alternate haplotypes are quite different in sequence. Comparison of the larger genomic regions containing these loci in each isolate (bin 06 in *Pgt*21-0 and bins 15 and 23 in Ug99) indicated that one haplotype (designated A) was >99.5% identical in Ug99 and *Pgt*21-0 (Fig. 2b, Supplementary Fig. 2 and Table 6). The other two haplotypes (B and C) were highly divergent from each other and from haplotype A, sharing only 62-75% total identity with many large insertions and deletions. The high similarity between the A haplotypes of this chromosome suggested that Ug99 and *Pgt*21-0 may share large portions of their genomes, potentially up to an entire haploid genome copy.

**Fig. 2.**
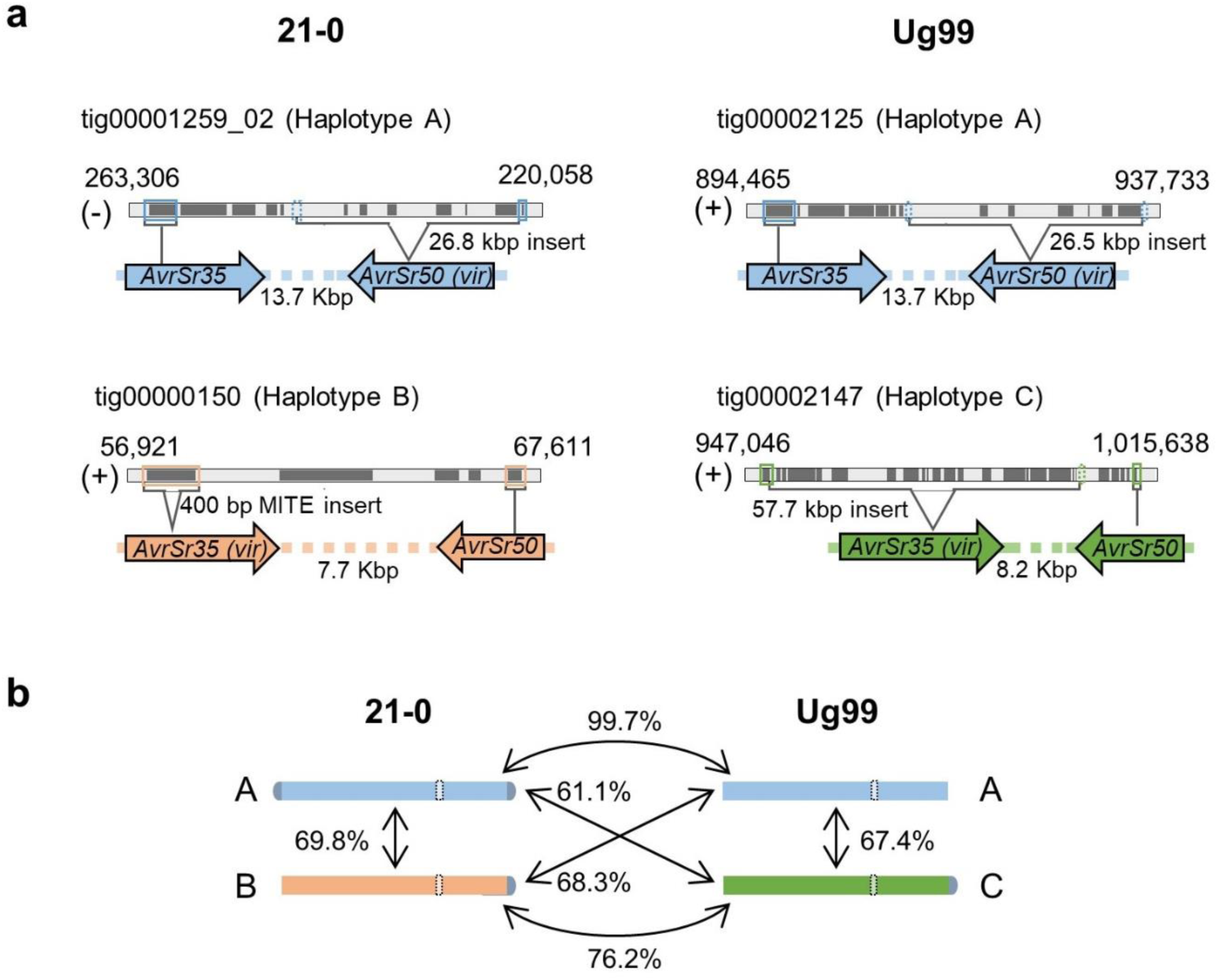
A common haplotype containing *AvrSr50* and *AvrSr35* is shared between *Pgt Pgt*21-0 and Ug99. **a,** Diagram of genomic regions containing *AvrSr50* and *AvrSr35* alleles in *Pgt*21-0 and Ug99. Numbers above tracks correspond to contig coordinates and the sense of the DNA strand is indicated as + or -. Predicted gene models are depicted as dark grey boxes and intergenic spaces are shown in light grey. *AvrSr50* and *AvrSr35* coding sequences are boxed and the direction of transcription is represented by coloured arrows, with intergenic distances indicated. Positions and sizes of insertions in virulence (*vir*) alleles are indicated by brackets. **b,** Total sequence identity between contigs representing homologous chromosomes of different haplotypes (coloured bars) containing the *AvrSr50*/*AvrSr35* locus (dotted white boxes). Telomere sequences are represented in grey. Chromosome size = ∼3.5 Mbp.

### Whole-genome haplotype assignment and comparison

Genome regions that shared high identity between Ug99 and *Pgt*21-0 were identified using a read subtraction and mapping approach (Fig. 3). Shared sequences were designated as haplotype A, while sequences unique to *Pgt*21-0 or Ug99 were designated as haplotypes B or C respectively (Supplementary Table 7). Some contigs in each assembly appeared to be chimeric with distinct regions assigned to opposite haplotypes, and these contigs were divided into separate fragments (Supplementary Table 8) for subsequent haplotype comparisons. Approximately half of each genome assembly was assigned to either the A, B or C haplotypes (Fig. 3c) and importantly one set of homologous sequences from each bin was assigned to each haplotype (Supplementary Table 8). The A, B and C haplotype sets contained 95-96% of conserved fungal genes (Fig. 3c), indicating that each represents a full haploid genome equivalent. Consistent with this, the haplotypes were highly contiguous (Fig. 3d). Overall sequence identity between the A haplotypes of *Pgt*21-0 and Ug99 was 99.5%, with structural variation (large insertions/deletions) representing only 0.5% of the haplotypes (Fig. 4a, Table 1, and Supplementary Table 9). In contrast, total sequence identity between the A, B or C haplotypes ranged between 87% and 91%, with structural variation accounting for 6.7% to 8.7% of the haploid genome sizes (Fig. 4b to d, Table 1 and Supplementary Table 9). There were only ∼9,000 SNPs (0.1/kbp) between the two A haplotypes, versus 876,000 to 1.4 million SNPs (11-18/kbp) between the A, B and C pairs, which is consistent with estimates of heterozygosity levels in *Pgt Pgt*21-0^18^. These similarities were supported by Illumina read coverage analysis (Supplementary Fig. 3), showing that Ug99 and *Pgt*21-0 share one nearly identical haploid genome copy.

**Fig. 3.**
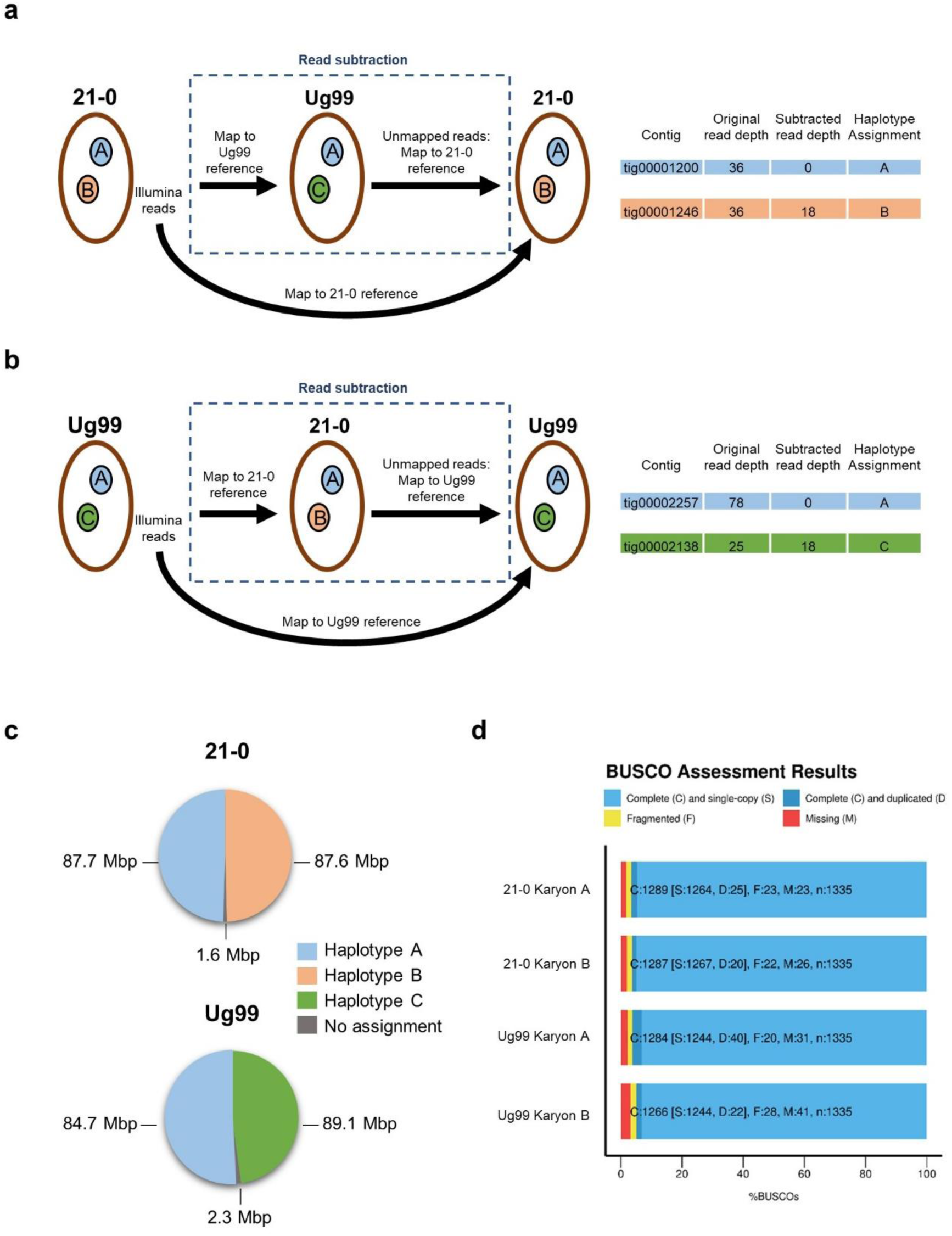
Haplotype assignment by read subtraction and mapping process. **a,** Illumina reads from *Pgt Pgt*21-0 were mapped to the Ug99 genome assembly at high stringency. Unmapped reads derived from divergent regions of the B haplotype were retained and then mapped to the *Pgt*21-0 genome assembly. Read coverage of individual contigs with the original and subtracted reads were compared to designate haplotypes as either A or B. **b,** The same process was followed with reads from Ug99 subtracted against the *Pgt*21-0 reference to designate the A and C haplotypes. **c,** Pie chart showing proportion and total sizes of contigs assigned to haplotypes A, B or C or unassigned in *Pgt Pgt*21-0 and Ug99 assemblies. **d,** BUSCO analysis to assess completeness of haplotype genome assemblies. Bars represent the percentage of total BUSCOs as depicted by the colour key.

**Fig. 4.**
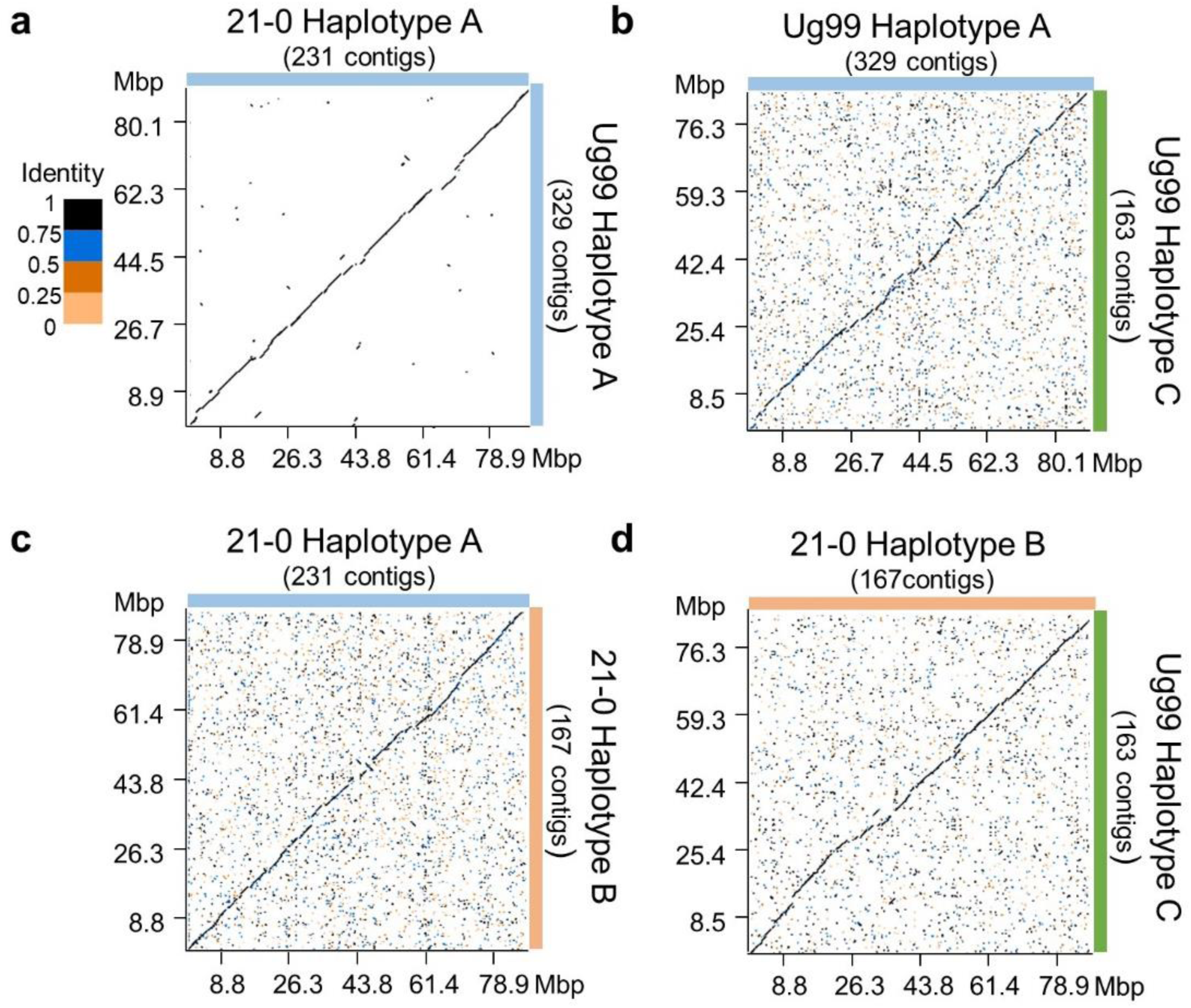
*Pgt Pgt*21-0 and Ug99 share one nearly identical haploid genome. **a** to **d,** Dot plots illustrating sequence alignment of complete haplotypes. X- and y-axes show cumulative size of the haplotype assemblies depicted by coloured bars to the right and top of the graphs. Colour key indicates sequence identity ratios for all dot plots.

**Table 1.** Intra- and inter-isolate sequence comparison of entire haplotypes in *Pgt* Ug99 and *Pgt*21-0.

| Sequence similarity |  |  | Structural variation |  |  |
| --- | --- | --- | --- | --- | --- |
| Isolate comparison | Bases aligned (%) | Sequence divergence (%) | Number of variants | Total variant size |  |
|  |  |  |  | Mbp | % of genome |
| 21-0 A vs Ug99 A | 99.64 | 0.08 | 491 | 0.82 | 0.46 |
| Ug99 A vs Ug99 C | 91.52 | 4.08 | 2,571 | 13.69 | 7.88 |
| 21-0 A vs *Pgt*21-0 B | 91.38 | 4.19 | 2,696 | 15.01 | 8.56 |
| 21-0 B vs Ug99 C | 93.44 | 2.4 | 1,910 | 11.50 | 6.69 |

### Assessment of inter-nuclear recombination and chromosome assembly

We tested two hypotheses that could explain the shared haplotype between Ug99 and *Pgt*21-0: 1) Ug99 arose by a somatic hybridisation event in which an isolate of the race 21 lineage donated an intact nucleus of the A haplotype (Fig. 5a); and 2) Ug99 arose by a sexual cross in which one haploid pycnial parent was derived from a race 21 lineage isolate after meiosis (Fig. 5b). Under both scenarios the A haplotype of Ug99 represents one entire haploid nucleus that was derived from the race 21 lineage isolate. In the nuclear exchange scenario, the *Pgt*21-0 A haplotype represents a single nucleus donated intact to generate Ug99. However, under the sexual cross model, this *Pgt*21-0 haplotype would include segments of both nuclear genomes that were combined by crossing over and chromosome reassortment after karyogamy and meiosis. Because the *Pgt*21-0 and Ug99 genome assemblies represent the phased dikaryotic state of each isolate, all correctly phased contigs in Ug99 should be either A or C haplotype, while those in *Pgt*21-0 would include mixed haplotype contigs only if the sexual cross hypothesis is correct. In fact, just 19 (out of 469) contigs in the Ug99 assembly appeared to be chimeric with adjacent regions of either the A or C haplotype. These cannot be explained biologically under either model, and appeared to result from haplotype phase swap artefacts. All the junctions occurred at positions corresponding to gaps between the corresponding alternate contigs, and Illumina read mapping showed that these sites contained either collapsed haplotype, non-unique sequences or discontinuities in read coverage (Supplementary Fig. 4), indicative of assembly errors disrupting phase information across the junction. Likewise, 31 contigs of mixed haplotype in the *Pgt*21-0 assembly all contained likely phase swap artefacts (Supplementary Fig. 4). To experimentally distinguish between phase-swap assembly artefacts and meiotic recombination events, we used Hi-C chromatin cross-linking proximity analysis ^26^ to assess physical linkage between binned contigs in the *Pgt*21-0 assembly. For each of the chimeric contigs, the separated A and B fragments showed significantly more Hi-C read pair connections to contigs of the same haplotype than to contigs of the other haplotype, including other fragments of the original chimeric contig (Supplementary Table 10). This analysis confirmed that all potential recombination sites in the *Pgt*21-0 genome assembly relative to Ug99 resulted from assembly phase swaps and not genetic recombination.

**Fig. 5.**
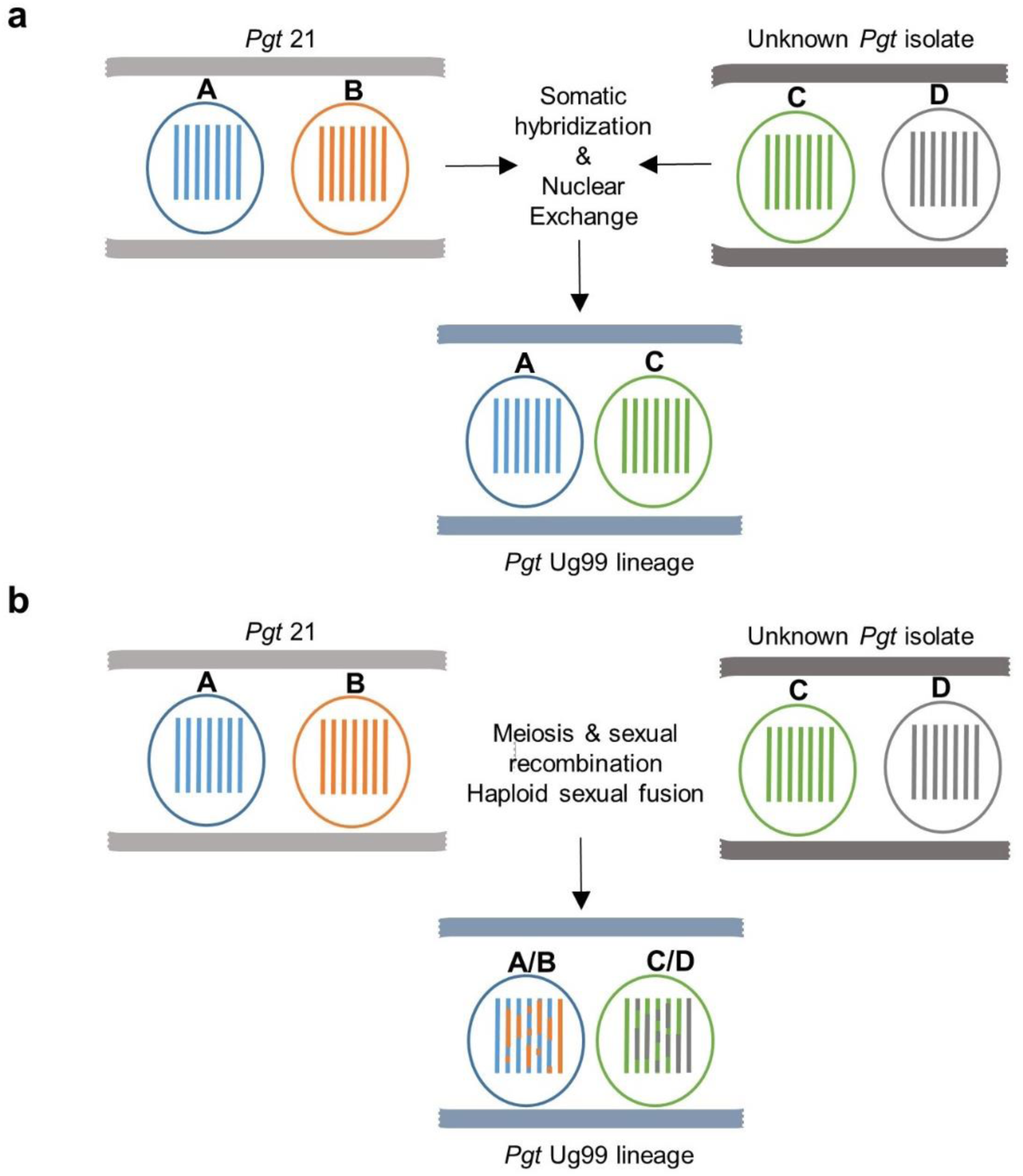
Models for the emergence of the founder isolate of the Pgt Ug99 lineage. **a,** A somatic hybridisation event and nuclear exchange occurred between an isolate of the Pgt 21 lineage and an unknown Pgt isolate. The combination of nuclei A and C yielded the parental isolate of the Ug99 lineage in Africa. Under this scenario, nucleus A of Ug99 is entirely derived from nucleus A in *Pgt*21-0. **b,** Alternatively, sexual reproduction and mating between these two parental isolates defined the origin of the Ug99 lineage. Under this scenario, meiotic recombination and chromosome reassortment would result in the *Pgt*21-0-derived A nucleus of Ug99 being composed of a mosaic of the two haploid nuclear genomes of *Pgt*21-0 (X and Y).

Combining Hi-C scaffolding data with the Bin and haplotype assignment information for the *Pgt*21-0 assembly allowed us to construct 18 chromosome pseudomolecules for each of the A and B haplotypes (Fig. 6a, Supplementary Table 11 and S12). These covered a total of 170 Mbp and ranged from 2.8 to 7.3 Mbp in size, consistent with relative chromosome sizes from karyotype analysis ^23^. Comparison of the A and B chromosomes showed high collinearity but detected two translocation events (Supplementary Fig. 7). These were supported by contigs that spanned the translocation breakpoints and by Hi-C linkages across these junctions. Approximately 65% of the total Hi-C read pairs represented links between physically contiguous sequences on the same chromosome, while the remaining pairs connected sites distributed across the genome. Because Hi-C DNA crosslinking is performed in intact cells, these non-scaffolding linkages should preferentially form between chromosomes that are located in the same nucleus. Indeed, all chromosomes of the A haplotype showed a much higher proportion of Hi-C read pair links to other chromosomes of the A haplotype (∼85%) than to chromosomes of the B haplotype (∼15%) (Fig. 6b), suggesting that they are all located in the same nucleus. Similarly, 17 of the B haplotype chromosomes showed stronger linkage to other B chromosomes (∼90%) than to A chromosomes (∼10%) (Fig. 6c). However, chromosome 11B showed the inverse, suggesting that both homologs of this chromosome are located in the same nucleus and implying a chromosome exchange event during asexual propagation of the *Pgt*21-0 isolate, after the exchange event leading to Ug99.

**Fig. 6.**
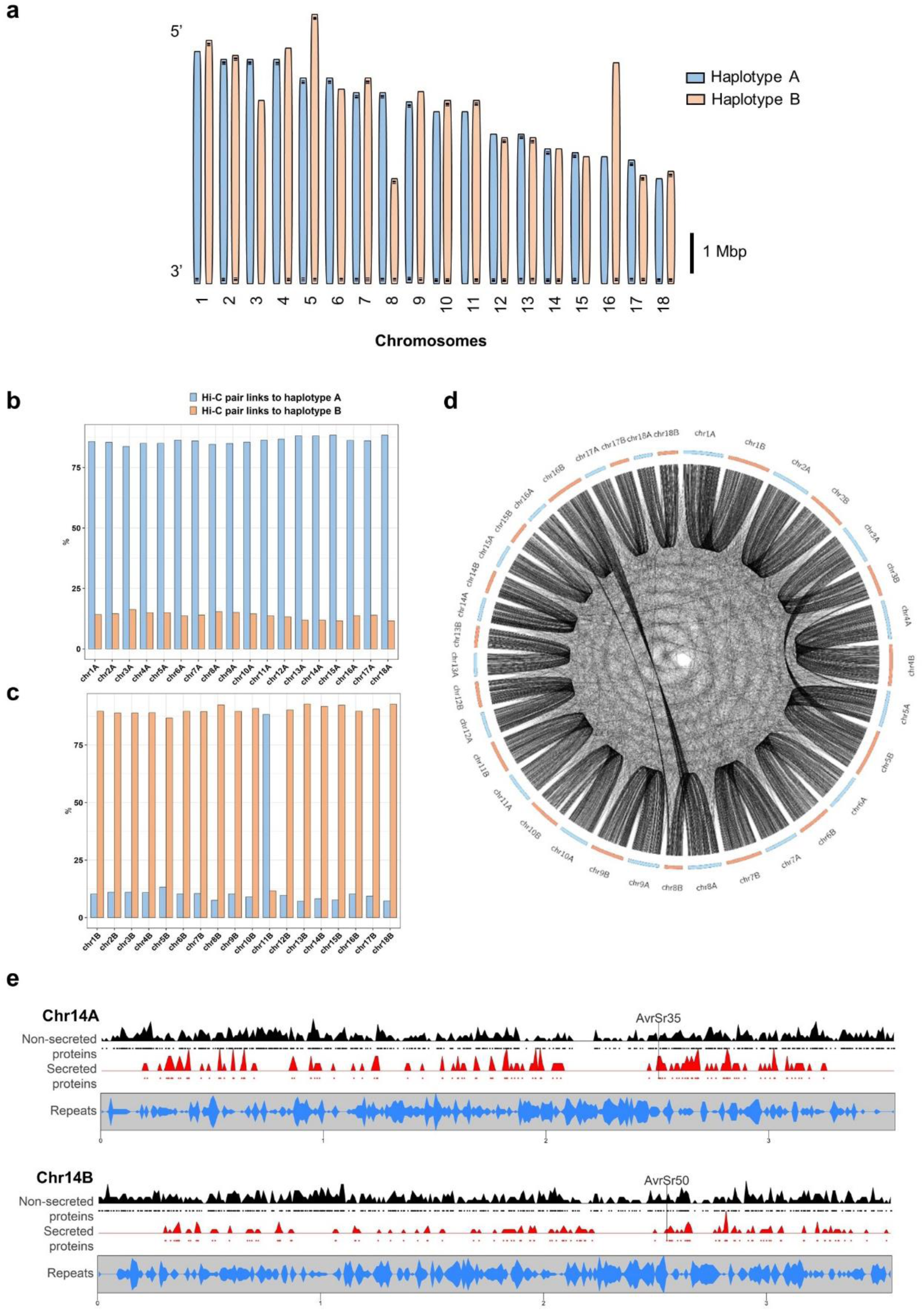
Chromosome sets of haplotype A and B in *Pgt Pgt*21-0. **a** Schematic representation of assembled chromosomes for *Pgt Pgt*21-0 of each haplotype (scale bar = 1 Mbp). Vertical bars indicate telomeric repeat sequences. **b** Percentage of Hi-C read pairs linking each A haplotype chromosome to other A chromosomes A (blue) or to B haplotype chromosomes (orange). **c** Percentage of Hi-C read pairs linking each B haplotype chromosome to either A (blue) or B (orange) chromosomes. **d** Gene and repeat density plots for homologous chromosomes 14A and 14B. Density of genes encoding non-secreted (black) or secreted proteins (red) along the chromosomes are shown, with individual genes indicated by black or red dots. Bottom graph shows density of repeat elements (blue). Positions of *AvrSr50* and *AvrSr35* genes are indicated. **e** Circos plot showing location of othologous gene pairs in the A and B chromosomes of *Pgt*21-0. Orthologous pairs are connected by black lines. This illustrates high level of synteny between haplotypes. Reciprocal translocations between chromosomes 3 and 5 as well as between chromosomes 8 and 16 can be observed.

Overall the whole genome comparison data demonstrate that Ug99 shares one full haploid nuclear genome with the *Pgt*21-0 isolate with no recombination events within chromosomes and no reassortment of chromosomes from different nuclei. These facts are inconsistent with a sexual origin, and strongly support that the Ug99 lineage arose by a somatic hybridisation event involving one parent derived from the African race 21 lineage and another parent of unknown origin (Fig. 7).

**Fig. 7.**
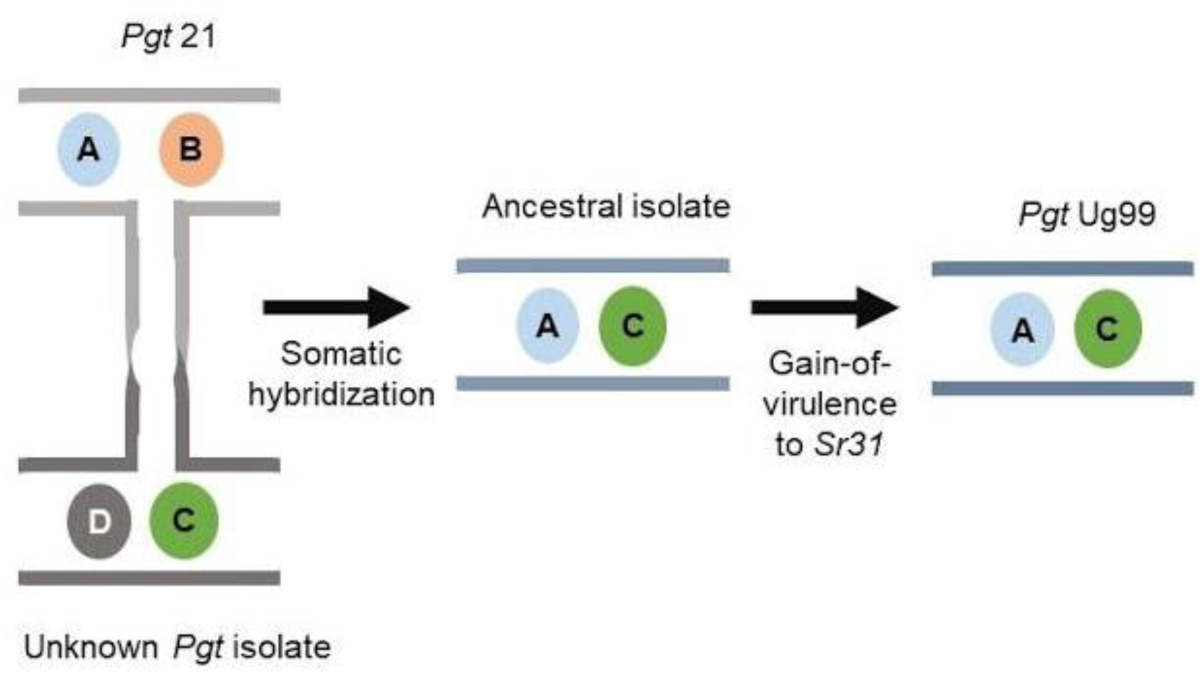
Model for Ug99 origin by somatic hybridisation and nuclear exchange between an isolate of the *Pgt* 21 lineage and an unknown *Pgt* isolate. The ancestral isolate of the lineage acquired the A and C genomes and later gained virulence to wheat cultivars carrying the *Sr31*resistance gene.

**Fig. 8.**
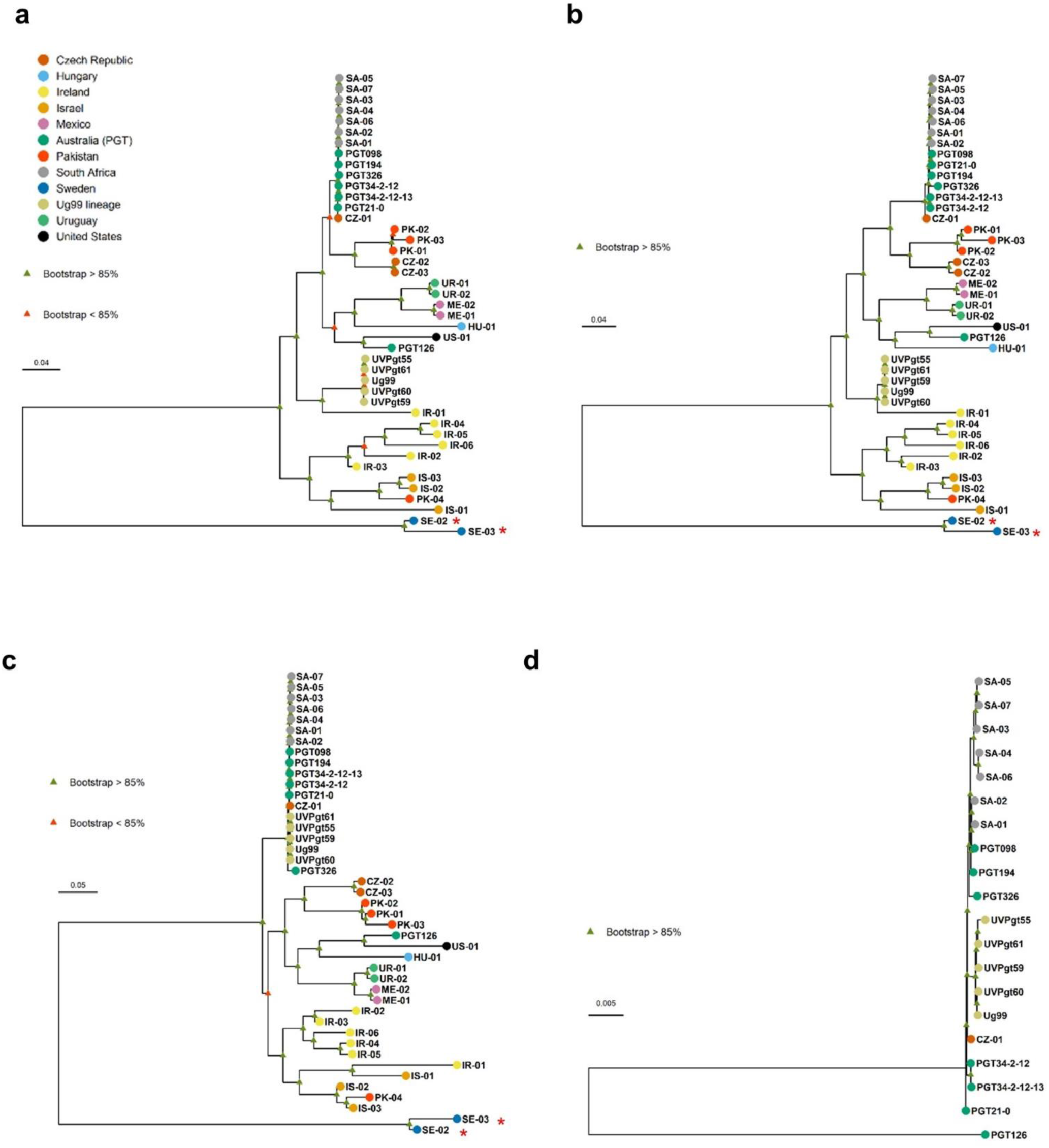
Somatic hybridisation in *Pgt* evolution. **a,** Phylogenetic analysis of *Pgt* isolates from diverse countries of origin (colour key) using a RAxML model and SNPs called against the full dikaryotic genome of *Pgt Pgt*21-0. Scale bar indicates number of nucleotide substitutions per site. Red asterisks indicate *P. graminis* f. sp. *avenae* isolates used as outgroup. **b,** Dendrogram inferred using biallelic SNPs detected against haplotype A of *Pgt Pgt*21-0. **c,** Dendrogram inferred using SNPs detected against haplotype B of *Pgt Pgt*21-0. **d,** Dendrogram inferred from SNPs detected in haplotype C of Ug99.

To compare gene content between defined haploid genomes of *Pgt*, we annotated the *Pgt*21-0 and Ug99 genome assemblies. Similar gene numbers were identified in each isolate, roughly equally distributed between the haplotypes (Supplementary Table 13). Gene orthology analysis indicated that 65-70% of genes in each of the A, B and C haplotypes were shared and represent a core *Pgt* gene set, while the remainder were present in only one or two haplotypes (Supplementary Table 14). Mapping of orthologous gene pairs supported the synteny of the *Pgt*21-0 chromosome assemblies (Fig. 6d). Genes encoding secreted and non-secreted proteins were similarly distributed across the chromosomes and showed an opposite distribution to repeat sequences (Fig. 6e Supplementary Fig. 5), consistent with the absence of two-speed genome architecture in rust fungi^12,13^. Both Ug99 and *Pgt*21-0 are heterozygous at the predicted *a* and *b* mating type loci (Supplementary Fig. 6) consistent with an expected requirement for formation of a stable dikaryon^11^.

### Phylogenetic analysis of global *Pgt* isolates

We used the haplotype-phased genome references for *Pgt*21-0 and Ug99 to determine genetic relationships within a set of global *Pgt* isolates using publicly available sequence data^18,24,27^. Maximum likelihood trees based on whole genome SNPs (Fig. 7a and Supplementary Fig. 7) showed a very similar overall topology to that reported previously for most of these isolates^27^. The five isolates of the Ug99 lineage, and the thirteen South African and Australian isolates each formed a separate tight clade, consistent with their proposed clonal nature^19-21^. However, tree building using filtered SNPs from just the A haplotype resulted in the Ug99, South African and Australian isolates forming a single clade, indicating the clonal derivation of this nucleus among these isolates (Fig. 7b, Supplementary Fig. 7). In contrast these groups remained in two distant clades in phylogenies inferred using filtered SNPs in the B genome. However, in this case two isolates from the Czech Republic and three isolates from Pakistan were now located in a single clade with the South African and Australian isolates (Fig. 7c). This suggests that these isolates contain a haplotype closely related to the B genome of the race 21 lineage and may also have arisen by somatic hybridisation and nuclear exchange. A phylogeny based on the C genome SNPs grouped isolate IR-01 from Iran with the Ug99 lineage (Fig. 7d), suggesting that these isolates share the C haplotype. IR-01 could represent a member of the parental lineage that donated the C nucleus to Ug99, or alternatively may have acquired the C nucleus from Ug99. Notably, this was the only isolate that shared the *AvrSr35* 57kbp insertion allele identified in Ug99 (Supplementary Fig. 8). The relationships between these putative hybrid isolates were also supported by the patterns of homozygous and heterozygous SNPs detected in each haplotype (Supplementary Fig. 8). The incongruities between phylogenies generated based on different haplotypes highlights the difficulty of inferring relationships between isolates based on whole genome SNP data without haplotype resolution. Overall, these observations suggest that somatic hybridisation and nuclear exchange may be a common mechanism generating genetic diversity in global populations of *Pgt*.

## Discussion

Although sexual reproduction of *Pgt* can generate individuals with novel genetic combinations, the completion of the sexual cycle requires infection of an alternate host, common barberry. In parts of the world where barberry is scarce or absent, either due to eradication programs or its natural distribution^28^, *Pgt* is restricted to asexual propagation with new diversity arising by mutation^19,21^. Somatic hybridisation provides an alternative explanation for the appearance of new races not derived by stepwise mutation. Hybrids with high adaptive value in agroecosystems may establish new lineages of epidemiological significance, as shown by the emergence of the Ug99 lineage with its substantial impact on East African wheat production and threat to global food security^14,29^. The role of somatic exchange in population diversity of other rust fungi is not known, although genetic exchange in experimental settings has been reported for several species^5,8,9,30^. Our findings provide a new framework to take advantage of haplotype resolution to understand population biology of rust fungi.

Extended dikaryotic developmental stages are common in many other fungi, especially basidiomycetes. Indeed, separation of karyogamy (fusion of haploid nuclei to form a diploid nucleus) from gamete fusion is a feature unique to the fungal kingdom^1^. However, it is unclear why fungi maintain an extended dikaryotic stage prior to formation of a diploid nucleus as a precursor to sexual reproduction ^31^. One possibility is that the ability to exchange haploid nuclei offers an advantage over the diploid state due to the enhanced genetic variation in long-lived asexual dikaryotes. Although there is now clear evidence of nuclear exchange between dikaryons, nothing is known of how this process occurs or is regulated. It differs from parasexuality in ascomycetes^2^, as the dikaryotic state is maintained with no nuclear fusion or haploidization Wang and McCallum^32^ observed the formation of fusion bodies where germ tubes of different *P. triticina* isolates came into contact, with the potential for nuclear exchange at these junctions. The possibility of genetic exchange between haploid nuclei in rust has also been proposed^6^, and in support of this we saw evidence for translocation of one chromosome between nuclei in *Pgt*21-0. There is also evidence for somatic exchange of genetic markers in dikaryotes of the mushroom *Schizophyllum commune*, which belongs to another Basidiomycete subphylum, Agaricomycotina^33^. Arbuscular mycorrhizae (AM) fungi (Mucoromycetes) are another ancient fungal lineage whose spores contain hundreds of nuclei, and for which no sexual stages have been described raising questions of how these lineages have survived^34^. Recently some dikaryotic-like AM isolates possessing two divergent classes of nuclei have been observed. Nuclear exchange between dikaryotes and/or between nuclei could be another driver of genetic variation in these fungi. Evidently, the members of the fungal kingdom display remarkable genetic plasticity and further investigation is required to reveal the mechanism, prevalence, and evolutionary importance of nuclear exchange in dikaryotic and multinucleate fungi.

## Materials and Methods

### Fungal stocks and plant inoculation procedures

*Puccinia graminis* f. sp. *tritici* (*Pgt*) isolates Ug99^15^, UVPgt55, UVPgt59, UVPgt60 and UVPgt61 collected in South Africa^16,35^ were transferred to the Biosafety Level 3 (BSL-3) containment facility at the University of Minnesota for growth and manipulation. Samples were purified by single pustule isolation and then amplified by 2-3 rounds of inoculation on the susceptible wheat cultivar McNair. Virulence pathotypes and purity of each isolate were confirmed by inoculation onto the standard wheat differential set (Supplementary Table 1) ^23,36^. An Australian isolate of pathotype 21-0 was first isolated in 1954 and has been described ^18,19^. North American isolate CRL 75-36-700-3 (pathotype SCCL) and Kenyan isolate 04KEN156/04 (pathotype TTKSK) were described previously^17,22^. For rust inoculations, urediniospores retrieved from −80 °C were activated by heat treatment at 45 °C for 15 min and suspended in mineral oil (Soltrol 170, Philips Petroleum, Borger, TX, U.S.A.) at 14 mg/ml. Seven day-old seedlings were spray-inoculated at 50 μl/plant and oil was allowed to evaporate. Inoculated plants were kept in a dark mist chamber at 22–25 °C with 100% humidity (30 min continuous misting followed by 16 h of 2 min misting at 15 min intervals). Subsequently, plants were exposed to light (400 W sodium vapor lamps providing 150–250 μmol photons s^-1^ m^-2^) for 3.5 h of 2 min misting at 15 min intervals and 2 h of no misting. After plants were dry, plants were transferred to a growth chamber under controlled conditions (18 h/6 h of light/dark, 24 °C/18 °C for day/night, 50% relative humidity). Spores were collected and maintained at −80 °C at 9 days post inoculation (9 dpi) and 14 dpi.

### DNA extraction and sequencing of rust isolates

High molecular weight DNA of Ug99 and *Pgt*21-0 was extracted from 300-350 mg urediniospores as previously described^37^, with the following modifications: 1) for Phenol:Chloroform:Isoamyl alcohol extractions, samples were centrifuged at 4 °C and 5,000 x g for 20 mins instead of 6,000 x g for 10 min; 2) a wide-bore 1mL pipette tip was used to transfer the DNA pellet; 3) samples were incubated for 1 h at 28°C with 200-250 rpm shaking to dissolve the final DNA pellet. Double stranded DNA concentration was quantified using a broad-range assay in a Qubit Fluorometer (Invitrogen, Carlsbad, CA, U.S.A.) and a NanoDrop (Thermo Fisher Scientific, Waltham, MA, U.S.A.). Approximately 10 µg DNA from Ug99 and *Pgt*21-0 was sequenced using PacBio single-molecule real-time (SMRT) sequencing (Pacific Bioscience, Menlo Park, CA, U.S.A.) at either the Frederick National Laboratory for Cancer Research, Leidos Biomedical Research, Inc. (Frederick, MD, U.S.A.) or the Ramaciotti Centre (Sydney, Australia) respectively. DNA was concentrated and cleaned using AMPure PB beads for Ug99 or AMPure XP beads for *Pgt*21-0 (Pacific Biosciences, Menlo Park, CA, U.S.A.). DNA quantification and size assessment was conducted using a NanoDrop (Thermo Fisher Scientific, Waltham, MA, U.S.A.) and 2200 TapeStation instruments (Agilent Technologies, Santa Clara, CA, U.S.A.). DNA was sheared to a targeted average size of 20 kb using G-tubes (Covaris, Woburn, MA, U.S.A.) according to the manufacturer’s instructions. Libraries were constructed following the 20 kb Template Preparation BluePippin Size-Selection System protocol (Pacific Biosciences) using a BluePippin instrument (Sage Science, Beverly, MA, U.S.A.) with a 0.75% agarose cassette and a lower cutoff of 15 kbp (protocol “0.75% agarose DF Marker S1 High Pass 15-20 kb”). For *Pgt* Ug99, 5 SMRT cells were sequenced on a PacBio Sequel platform using P6-C4 chemistry, the Sequel Binding Kit 2.0 (Pacific Biosciences), diffusion loading, 10-hour movie lengths and Magbead loading at 2 pM (3 cells) or 4 pM (2 cells). In addition, 4 SMRT cells were run on PacBio RSII sequencer using P6-C4 chemistry (Pacific Biosciences), with 0.15 nM MagBead loading and 360-min movie lengths. For *Pgt Pgt*21-0, 17 SMRT cells were run on the RSII platform using P6-C4 chemistry, Magbead loading (0.12-0.18 nM) and 240-min movie lengths.

Genomic DNA for Illumina sequencing was extracted from 10-20 mg urediniospores of Ug99, UVPgt55, 59, 60 and 61 using the OmniPrep™ kit (G-Biosciences, St. Louis, MO, U.S.A.) following the manufacturer’s instructions. TruSeq Nano DNA libraries were prepared from 300 ng of DNA and 150bp paired-end sequence reads were generated at the University of Minnesota Genomics Center (UMGC) on the Illumina NextSeq 550 platform using Illumina Real-Time Analysis software version 1.18.64 for quality-scored base calling.

### *De novo* long read assembly of *Pgt* Ug99 and *Pgt*21-0

Genome assemblies of *Pgt* Ug99 and *Pgt*21-0 were built from PacBio reads using Canu version 1.6^38^ with default parameters and an estimated dikaryotic genome size of 170 Mbp. Assemblies were first polished using the raw PacBio reads with the Arrow variant-calling algorithm in the pre-defined resequencing pipeline (sa3_ds_resequencing) in pbsmrtpipe workflow within SMRTLINK/5.1.0 (Pacific BioSciences). Assemblies were further polished by two rounds of Pilon^39^ with the option fix --all using Illumina reads generated from Ug99 in this work or previously generated reads from *Pgt*21-0 (NCBI SRA runAccession# SRR6242031). A BLASTN search (version 2.7.1)^40^ against the NCBI nr/nt database (downloaded on 4/11/2018) with E-value set as 1e-10 identified two contigs in the Ug99 assembly with significant hits to plant rRNA and chloroplast sequences and these were removed. PacBio and Illumina reads were mapped to the assembly using BWA-MEM (version 0.7.17)^41^ and BAM files were indexed and sorted using SAMtools (version 1.9)^42^. Read coverage analysis using genomeCoverageBed in BEDtools (version 2.27.1)^43^ identified 144 small contigs (< 50 kbp) in the Ug99 assembly with low coverage (< 2X) for both short and long reads mapping and these contigs were also excluded from the final assembly. Genome assembly metrics were assayed using QUAST (version 4.3)^44^. Genome completeness was assessed via benchmarking universal single-copy orthologs (BUSCOs) of the basidiomycota as fungal lineage and *Ustilago maydis* as the species selected for AUGUSTUS gene prediction^45^ in the software BUSCO v2.0 (genome mode)^46^. Telomeric sequences were identified using either a high stringency BLAST with 32 repeats of TTAGGG as query or a custom python script to detect at least five CCCTAA or TTAGGG repeats in the assemblies (github: https://github.com/figueroalab/Pgt_genomes). Repeats of at least 60 bp length and occurring within 100 bp of the 5’ or 3’ ends of the contig were defined as telomeric sequences.

### Detection of alternate contigs and bin assignment

To identify contigs representing corresponding haplotypes we used a gene synteny based approach (Fig. 1). The 22,484 predicted *Pgt* gene coding sequences^18^ were screened against the genome assemblies using BLITZ (Blat-like local alignment) in the Biokanga Tool set, (https://github.com/csiro-crop-informatics/biokanga/releases/tag/v4.3.9). For each gene the two best hits in the assembly were recorded. In most cases these will correspond to the two allelic versions of the gene, one in each haplotype. Thus contigs sharing best hits for at least five genes were selected as potential haplotype pairs and their sequence collinearity was examined by alignment and similarity plotting using D-genies^47^. Contigs representing contiguous or syntenous haplotypes were grouped together as bins after manual inspection.

### Identification of *AvrSr50* and *AvrSr35* region and validation of a 57 kbp insert in *AvrSr35*

Contigs containing the *AvrSr50* and *AvrSr35* gene sequences were identified by BLASTN search against customized databases for the Ug99 and *Pgt*21-0 genome assemblies. Manual inspection of coordinates of the BLAST hits of *AvrSr35* in the Ug99 assembly identified one complete copy on tig00002125 and a second copy on tig00002147 that was interrupted by an insertion sequence of ∼57 kbp (97% and 99% identity for the two aligned 5’ and 3’ fragments). Illumina and PacBio reads of Ug99 mapped to the genome assembly were visualized in the Integrative Genomics Viewer (IGV) which verified the contiguity of reads across this insertion (Supplementary Fig. 1). To validate the presence of this insert, flanking and internal sequences of the 57 kbp insert in *AvrSr35* were amplified from genomic DNA extracted using the OmniPrep™ kit (G-Biosciences, St. Louis, MO, U.S.A.) from rust urediniospores of Ug99, 04KEN156/04, and CRL 75-36-700-3. PCR was performed using Phusion high-fidelity DNA polymerase according to the manufacturer’s recommendations (New England BioLabs Inc., Ipswich, MA, U.S.A.) and primers (Supplementary Table 15) designed using Primer3^48^. The amplified PCR products were separated by electrophoresis on a 1% agarose gel along with the GeneRuler 1 kb DNA ladder Plus (Thermo Fisher Scientific, Waltham, MA, U.S.A.) as marker. The gel was stained using SYBR Safe DNA gel stain (Invitrogen Life Technologies, Carlsbad, CA, U.S.A.) and specific bands were cleaned using NucleoSpin gel clean-up kit (Takara Bio, Mountain View, CA, U.S.A.) for subsequent Sanger sequencing at UMGC. Base calling was performed using Sequencher 4.10.1, and sequences were aligned using Clustal Omega^49^ to *AvrSr35* alleles extracted from the genome assembly. The diagram of predicted gene models in the *AvrSr35* and *AvrSr50* locus on the corresponding contigs was depicted based on gene prediction results in this study and a custom R script (github: https://github.com/figueroalab/Pgt_genomes) using GenomicFeatures^50^ and ggbio^51^.

### Haplotype assignment by read cross mapping and subtraction

We used a read subtraction and mapping approach (Fig. 3) to identify contigs in the *Pgt* Ug99 and *Pgt*21-0 assemblies that showed high similarity and may be derived from a shared haplotype. Illumina reads from each isolate were mapped at high stringency to the reference of the other isolate and those reads that failed to map were retained, thus subtracting out sequences that were common to both isolates. Coverage of the remaining subtracted reads on the original reference was then used to identify contigs representing either shared or isolate specific sequences. For this approach, Illumina reads from *Pgt*21-0 (NCBI SRR6242031) were trimmed (“Trim sequences” quality limit = 0.01) and mapped to the Ug99 reference assembly using the “map reads to reference” tool in CLC Genomics Workbench version 10.0.1 or later with high stringency parameters (similarity fraction 0.99, length fraction 0.98, global alignment). Unmapped reads (Ug99-subtracted reads) were retained and then mapped back to the *Pgt*21-0 assembly contigs using the same parameters. In this way reads derived from the shared A haplotype were selectively removed and reads from divergent regions of the B haplotype were retained. The original *Pgt Pgt*21-0 reads were also mapped to the *Pgt*21-0 assembly and the read coverage for each contig compared to the Ug99-subtracted reads. Contigs with very low coverage (<2X total and <10% of the original read coverage) with the Ug99-subtracted reads were designated as karyon A (Fig. 3, Supplementary Table 7). Contigs with substantial coverage of Ug99-subtracted reads (>20% of the original read coverage) were designated as karyon B. Contigs with ambiguous read mapping data, including those with low coverage in the original unsubtracted reads or covered by largely non-uniquely mapping reads were left as unassigned. Read mapping to all contigs was confirmed by visual inspection of coverage graphs and read alignments in the CLC Genomics Workbench browser. Potentially chimeric contigs were identified as containing distinct regions with either high or no coverage with the Ug99-subtracted reads (Supplementary Fig. 4). For subsequent comparison and analyses, these contigs were manually split into their component fragments which were designated as haplotype A or B accordingly (Supplementary Table 7). The same process was followed in reverse for the assignment of the A and C haplotype contigs in Ug99. Trimmed Ug99 Illumina reads were mapped to the *Pgt*21-0 reference and unmapped reads (21-0-subtracted) were retained for subsequent mapping to the Ug99 reference and comparison of read coverage with the original reads. In this case, contigs with low subtracted-read coverage were designated as haplotype A, while contigs with substantial retained coverage were designated as haplotype C. The completeness of haplotype assignment in *Pgt Pgt*21-0 and Ug99 was assessed using BUSCOs of the basidiomycota fungal lineage and *Ustilago maydis* as the species selected for AUGUSTUS gene prediction in the software BUSCO v2.0 (transcript and protein modes)^46^.

### Sequence comparisons of genome assemblies

Haplotype sequences of the *AvrSr50*/*AvrSr35* chromosome as well as the full haploid genomes were aligned using MUMmer4.x^52^, (https://github.com/mummer4/mummer/blob/master/MANUAL.md) with nucmer -maxmatch and other parameters set as default. The alignment metrics were summarized in the report files of MUMmer dnadiff. Structural variation between haplotypes was determined using Assemblytics^52^ from the MUMmer delta file with a minimum variant size of 50 bp, a maximum variant size of 100 kbp, and a unique sequence length for anchor filtering of 10 kbp. The haplotype alignments were visualized in dot plots using D-genies with default settings^47^.

### Read coverage analysis and SNP calling on haplotypes

Illumina reads from Ug99 and *Pgt*21-0 were each mapped against the Ug99 and *Pgt*21-0 assemblies in CLC Genomics Workbench (similarity fraction 0.98, length fraction 0.95). For each assembly the mean coverage per base was calculated per 1,000 bp interval (“window”) using samtools bedcov and read coverage frequency normalized to the mean coverage of each haplotype was graphed as a violin plot using seaborn 0.9.0 package (https://seaborn.pydata.org/) using a custom python script (github: https://github.com/figueroalab/Pgt_genomes). To detect SNPs between two haplotypes, Illumina read pairs of *Pgt Pgt*21-0 that mapped uniquely to either the *Pgt*21-0 A or B haplotype contigs were extracted. Similarly, Ug99-derived read pairs that uniquely mapped to either the A or C haplotype contigs of Ug99 were extracted. These read sets were then separately mapped to the two assemblies in CLC Genomics Workbench (similarity fraction 0.99, length fraction 0.98). Variant calling was performed using FreeBayes v.1.1.0^53^ with default parameters in parallel operation. High quality SNPs were called by vcffilter of VCFlib (v1.0.0-rc1, https://github.com/vcflib/vcflib) with the parameter -f “QUAL > 20 & QUAL / AO > 10 & SAF > 0 & SAR > 0 & RPR > 1 & RPL > 1”. Homozygous and heterozygous SNPs were extracted by vcffilter -f “AC > 0 & AC = 2” and -f “AC > 0 & AC = 1”, respectively. SNP statistics were calculated using vcfstats of VCFlib.

### Hi-C data analysis and scaffolding

A Hi-C library was constructed with the ProxiMeta Hi-C kit from Phase Genomics v 1.0 containing the enzyme Sau3A. About 150 mg of dried urediniospores of *Pgt Pgt*21-0 were used as starting material following the standard protocol with the following exceptions. Spores were washed in 1 mL 1x TBS buffer twice before cross-linking. After quenching of the crosslinking, all liquid was removed and the wet spores frozen in liquid nitrogen. Frozen spores were lysed using cryogenic bead beating with two 5 mm steel beads shaking twice for 45 sec at 25 Hz using TissueLyser II (Qiagen). Lysis buffer was added to the frozen broken spore pellet, vortexed until full suspension, and the standard protocol continued. Reverse cross-linking was performed at 65°C with 700 rpm horizontal shaking for 18 h. Afterwards the standard protocol was followed. The Hi-C library was sequenced (150 bp paired-end reads) on the NextSeq 550 System using the Mid-Output Kit at the Ramaciotti Centre (Sydney, Australia). The raw Hi-C reads were processed with the HiCUP pipeline version 0.7.1^54^ (maximum di-tag length 700, minimum di-tag length 100, --re1 ^GATC,Sau3A), using bowtie2 as the aligner^55^ and the *Pgt*21-0 genome assembly as the reference. HiCUP produces SAM files representing the filtered di-tags and these were parsed to extract cis-far pairs (pairs located on the same contig and >10 kbp apart) and trans pairs (located on different contigs). The numbers of trans pairs connecting each pair of contigs was extracted from this data.

For scaffolding, the raw Hi-C reads were first mapped to the *Pgt*21-0 assembly using BWA-MEM^56^. The Arima Genomics pipeline was followed to post-process the alignments and filter for experimental artifacts (https://github.com/ArimaGenomics/mapping_pipeline/blob/master/01_mapping_arima.sh). Then SALSA 2.2^57^ was run on the processed read alignments (-e GATC) to scaffold the assembly. SALSA scaffolding was performed independently on the full set of contigs, as well as on the two sets of contigs assigned as haplotype A or B (each including the contigs with no assigned haplotype). Invalid scaffold linkages between adjacent telomeres, which occur as an artefact of telomere co-location within the nucleus, were discarded. The three sets of scaffolds were compared with the Bin and haplotype assignment information to find overlaps, which resulted in a final grouping of 18 chromosome builds of haplotype A and 18 chromosomes of haplotype B. Chromosome pseudomolecules were constructed by concatenating ordered contigs with 100 Ns inserted between contigs. Two translocation events were detected in the A and B chromosome sets (Supplementary Fig. 7), one between chromosomes 3 and 5 and one between chromosomes 8 and 16. These were supported by contigs that spanned the translocation junctions in both haplotypes. To further confirm these translocations, these contigs were separated into two fragments at the junction point and the SALSA scaffolding process was repeated on the full genome contig assembly. In each case the original contig containing the translocation junction was re-assembled in the subsequent scaffolds, supporting that the original contig assembly was correct and represented true translocation events within the A or B genomes. To detect nucleus-specific cross-links between chromosomes, HiCUP analysis was performed using the chromosome pseudomolecules as the reference assembly and the proportion of trans linkages between chromosomes of the same or different haplotype computed.

### Gene prediction and functional annotation

The genome assemblies of *Pgt* Ug99 and *Pgt*21-0 (as chromosome pseudomolecules for *Pgt*21-0) were annotated using the Funannotate pipeline^58^ (https://github.com/nextgenusfs/funannotate). Contigs were sorted by length (longest to shortest) and repetitive elements were soft-masked using RepeatModeler (v1.0.11) and RepeatMasker (v4.0.5) with RepBase library (v. 23.09)^59,60^. RNAseq libraries from *Pgt Pgt*21-0 (Supplementary Table 16)^18,24^ were used for training gene models. In the training step, RNA-seq data were aligned to the genome assembly with HISAT2^61^. Transcripts were reconstructed with Stringtie (v1.3.4d)^62^. Genome-guided Trinity assembly (v2.4.0)^63^ and PASA assembly (v2.3.3)^64^ were performed. To assist in predicting effector-like genes, stringtie-aligned transcripts were used in CodingQuarry Pathogen Mode (v2.0)^65^. The prediction step of funannotate pipeline (funannotate predict) was run with --ploidy 2, --busco_db basidiomycota and default parameters. Transcript evidence included Trinity transcripts, Pucciniamycotina EST clusters downloaded and concatenated from JGI MycoCosm website (http://genome.jgi.doe.gov/pucciniomycotina/pucciniomycotina.info.html, April 24, 2017), and predicted transcript sequences of haustorial secreted proteins^18^. Transcript evidence was aligned to the genome using minimap2 v2.1.0^66^ and the protein evidence was aligned to genome via Diamond (v0.9.13)/Exonerate (v2.4.0)^67^ using the default UniProtKb/SwissProt curated protein database from funannotate. *Ab initio* gene predictor AUGUSTUS v3.2.3^45^ was trained using PASA data and GeneMark-ES v4.32^68^ was self-trained using the genome assembly. Evidence Modeler was used to combine all the transcript evidence and protein evidence described above, gene model predictions from AUGUSTUS and GeneMark-ES, PASA GFF3 annotations and CodingQuarry Pathogen Mode (CodingQuarry_PM) GFF3 annotations using default weight settings except that the weight of PASA and CodingQuarry_PM were both set to 20. tRNA genes were predicted using tRNAscan-SE v1.3.1^69^. Gene models including UTRs and alternative spliced transcripts were updated using RNAseq data based on Annotation Comparisons and Annotation Updates in PASA. Funannotate fix was run to validate gene models and NCBI submission requirements. Genome annotation was assessed using BUSCOs of the basidiomycota fungal lineage and *Ustilago maydis* as the species selected for AUGUSTUS gene prediction in the software BUSCO v2.0 (transcript and protein modes)^46^.

Functional annotation was performed using funannotate annotate. Protein coding gene models were firstly parsed using InterProScan5 (v5.23-62.0) which was run locally to identify InterPro terms, GO ontology and fungal transcription factors^70^. Pfam domains were identified using PFAM v. 32.0, and carbohydrate hydrolyzing enzymatic domains (CAZYmes) were annotated using dbCAN v7.0^71^. UniProt DB v 2018_11, MEROPS v. 12.0 were used for functional annotation using Diamond blastp^72-74^. BUSCO groups were annotated with Basidiomycota models, eggNOG terms were identified using eggNOG-mapper v1.0.3^75^.

Gene and repeat density plots for chromosomes were generated using karyoploteR^76^. A protein was labelled as secreted if it was predicted to be secreted by the neural network predictor of SignalP 3.0^77^ and if it had no predicted transmembrane domain outside the first 60 amino acids using TMHMM^78^. RepeatMasker 4.0.6 with the species fungi^60^ was used to softmask repeats. Repeats longer than 200 bp were used in the chromosome plotting.

### Detection of mating loci in *Pgt Pgt*21-0 and Ug99

Putative mating-type loci in *Pgt Pgt*21-0 and Ug99 were identified by BLAST search with the alleles of the pheromone peptide encoding genes (*mfa2* or *mfa3*) and pheromone mating factor receptors (*STE3.2* and *STE3.3*) from the *a* locus, and the divergently transcribed *bW*/*bE* transcription factors from the *b* locus that were previously identified in *Pgt* isolate CRL 75-36-700-3^79^. Based on the genome coordinates of the BLAST hits, the predicted mating-type genes were extracted from the Ug99 and *Pgt*21-0 genome annotation. Protein sequences were aligned in Clustal Omega^49^.

### Phylogenetic analysis of rust isolates

For whole genome SNP calling and phylogenetic analysis we used Illumina DNA sequence data (Supplementary Table 17) from the five Ug99 lineage isolates described here, seven Australian isolates we described previously^18,24^ as well as 31 global isolates ^27^ downloaded from the European Nucleotide Archive (ENA; PRJEB22223). All sequence data files were checked for read quality using FASTQC software^80^. Reads were trimmed with Trimmomatic Version 0.33^81^ using default settings for adaptor trimming and for base quality filtering and reads < 80 bp were discarded. Quality filtered reads were aligned to the *Pgt* Ug99 or *Pgt*21-0 genome assemblies using BWA program version 0.7.17^56^ and technical replicates were merged using SAMtools 1.6^42^ and PICARD toolkit (Broad Institute 2018, http://broadinstitute.github.io/picard/) to generate final sequence alignment map (SAM) files for downstream analysis. Read lengths and coverage were verified by the functions *bamtobed* and *coverage* in BEDtools^43^ and *flagstat* in SAMtools. Variants were detected using FreeBayes version 1.1.0^53^ to call biallelic SNP variants across the 43 samples simultaneously. VCF files were subjected to hard filtering using vcffilter in vcflib (v1.0.0-rc1)^82^ with the parameters *QUAL > 20 & QUAL / AO > 10 & SAF > 0 & SAR > 0 & RPR > 1 & RPL > 1 & AC > 0* to generate final VCF files for phylogenetic analysis. To verify that each sample consisted of a single genotype free of contamination, read allele frequencies at heterozygous positions^83^ were examined using the vcfR package^84^. VCF files were converted to multiple sequence alignment in PHYLIP format using the *vcf2phylip* script^85^ and *R*-package *ips/phyloch* wrappings^86^. Phylogenetic trees were constructed using the maximum likelihood criterion (ML) in RAxML version 8.2.1.pthread^87^, assuming unlinked loci and support for groups was assessed using 500 bootstrap replicates and a general time reversible (GTR) model. Convergence and posterior bootstopping (bootstrapping and convergence criterion) were confirmed with the *-I* parameter in RAxML and also with *R*-packages *ape*^88^, *ips/phyloch*^86^, and *phangorn*^89^. Trees were drawn using *ggplot2*^90^ and *ggbio*^51^ *R*-packages.

SNPs representing the A, B, or C haplotypes were separated from the total SNP sets based on bed files of the contig coordinates on each haplotype (Supplementary Table 8) using the function *intersect – header* in BEDtools. The frequency of homozygous and heterozygous SNPs for haplotype-separated SNP sets was counted using vcfkeepsamples and vcffixup. Homozygous and heterozygous SNPs were extracted by vcffilter -f “TYPE = snp” and -f “AC > 0 & AC = 2” and -f “AC > 0 & AC = 1”, respectively. SNP statistics were calculated using vcfstats of VCFlib (v1.0.0-rc1).

### Orthology analysis

Gene annotations with multiple isoforms were reduced to a representative isoform by selecting the longest CDS using a custom perl script. Orthologous proteins were identified with Orthofinder^91^ using default parameters. Multiple pairwise orthology analyses were run based on within-isolate and cross-isolate comparisons of similar haplotypes (i.e. A versus A or B versus C). Additional comparisons were made between *Pgt*21-0 A, *Pgt*21-0 B, and Ug99 C haplotypes, as well as between Ug99 A, Ug99 C, and *Pgt*21-0 B haplotypes.

### Data availability

Sequence data and assemblies described here are available in NCBI BioProjects XXXX. Assemblies and annotations will also be available at the DOE-JGI Mycocosm Portal. Unless specified otherwise, all scripts and files will be available at https://github.com/figueroalab/Pgt_genomes.

## Acknowledgments

We thank P. van Esse, G. Bakkeren, C. Aime and Y. Jin for valuable discussions, S. Dahl and N. Prenevost for technical support, J. Palmer for gene annotation troubleshooting, and the Minnesota Supercomputing Institute for computational resources. This research was funded by two independent grants from the TwoBlades foundation to P.N.D. and M.F., respectively, by a USDA-Agriculture and Food Research Initiative (AFRI) Competitive Grant (Proposal No. 2017-08221) to M.F, and University of Minnesota Lieberman-Okinow and Stakman Endowments to B.J.S. M.F. and M.E.M were supported by the University of Minnesota Experimental Station USDA-NIFA Hatch/Figueroa project MIN-22-G19 and an USDA-NIFA Postdoctoral Fellowship award (2017-67012-26117), respectively. B.S. is supported by an ARC Future Fellowship (FT180100024).

## Author contributions

M.F and P.N.D conceptualized the project, acquired funding and supervised the work. B.V. and Z.A.P. provided study materials. F.L., N.M.U., C.R., O.M., B.S., R.M., and B.J.S. acquired experimental data. F.L., N.M.U., J.S., B.S., B.J.S., H.N.P., P.N.D, K.S., E.H., M.E.M., and C.D.H. conducted data analysis. M.F. and P.N.D. drafted the manuscript. All authors contributed to review and editing.

---

**Supplementary Fig. 1.**
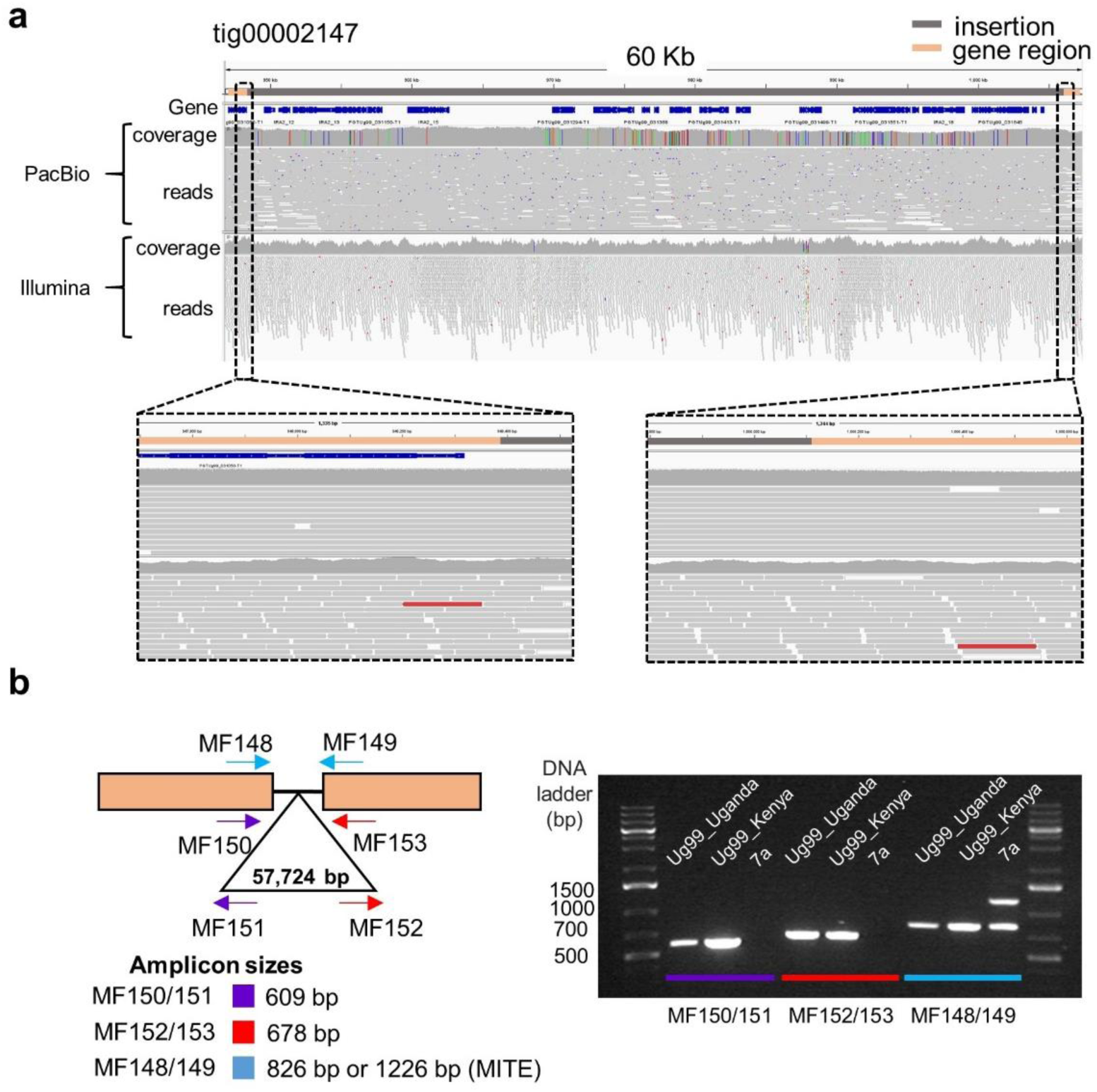
Presence of a 57 kbp-insert in one allele of *AvrSr35* in Ug99. **a,** Genome browser view of a 60 kbp genomic region in haplotype C of Ug99. The top bar shows the *AvrSr35* coding sequences (orange) flanking a 57 kbp-insert (grey). Annotated gene models (blue) are shown below. The following tracks show the read coverage graph and the alignments of Ug99 reads mapped to this region. Zoomed-in areas (boxed) show read mapping across the junction between the *AvrSr35* coding sequence and the 5’ and 3’ ends of the inserted sequence. **b,** Validation of 57 kbp-insert in *AvrSr35* of Ug99 isolates via PCR amplification. The positions of primers on the *AvrSr35* gene (orange boxes) and insertion (triangle) are shown along with the predicted amplicon sizes. PCR amplification products from the original Ug99 isolate (Ug99_Uganda), the Kenyan Ug99 isolate 04KEN156/04 (Ug99_Kenya)^24,25^ and the isolate CRL 75-36-700 (7a)^22^. Note that 7a is heterozygous for a wildtype allele of *AvrSr35* and a virulence allele containing a 400bp MITE insertion^25^.

**Supplementary Fig. 2.**
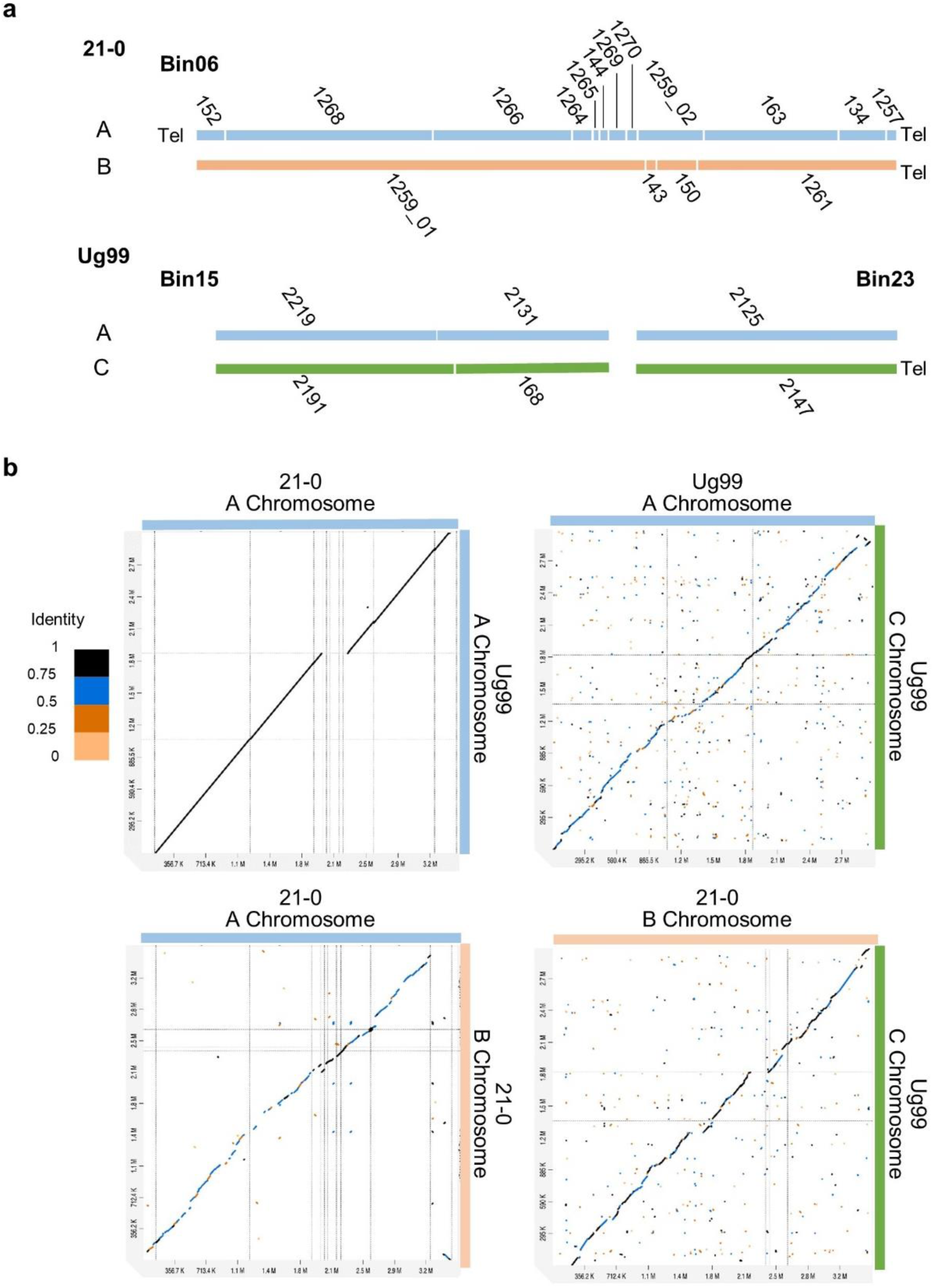
One of the homologous chromosomes containing *AvrSr50* and *AvrSr35* loci is nearly identical in *Pgt Pgt*21-0 and Ug99. **a,** Schematic representation of the alignment of contigs in *Pgt Pgt*21-0 and Ug99 derived from the homologous chromosomes. Contig IDs are indicated as numbers and presence of telomeres as “Tel”. The *Pgt*21-0 contigs were assembled as Bin06 and contain telomeres at both ends indicating that a full chromosome was represented. The homologous contigs from Ug99 were present in two bins (Bin15 and Bin23). Contigs are coloured according to haplotype designation; A (light blue); B (orange); C (green). **b**, Dot plots of alignments between the homologous chromosomes of each haplotype, indicated by coloured bars at the top and right. X- and y-axes represent nucleotide positions. Colour key indicates sequence identity fraction for all dot plots (1= maximum identity score).

**Supplementary Fig. 3.**
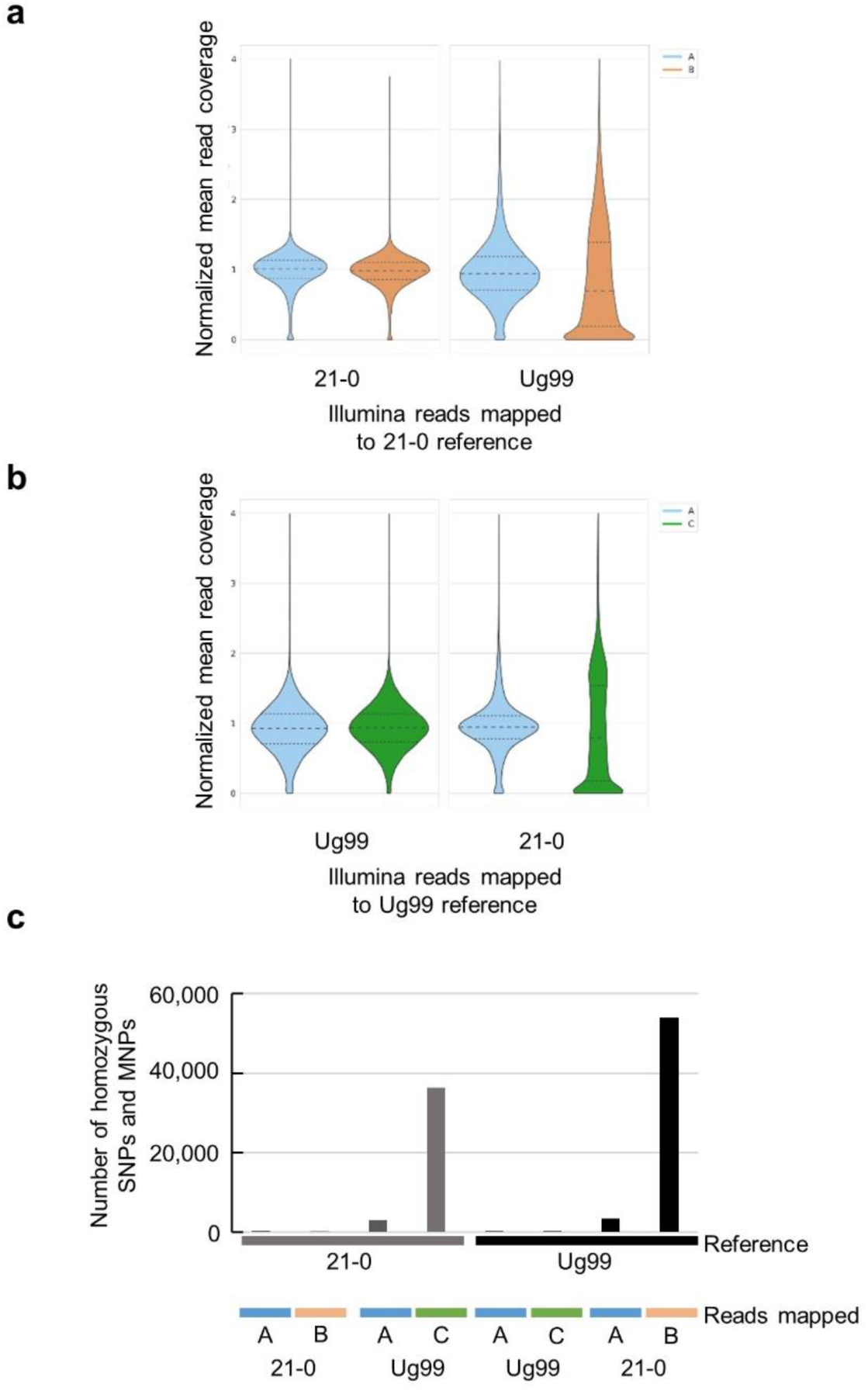
Haplotype-specific read mapping and SNP calling validates the close identity of haplotype A in *Pgt* Ug99 and *Pgt*21-0. **a,** Violin plots for the distribution of read coverage for haplotype A (blue) and B (orange) after mapping Illumina reads from *Pgt* Ug99 or *Pgt*21-0 to the *Pgt*21-0 assembly. **b**, Violin plots for the distribution of read coverage for haplotype A (blue) and C (green) after mapping Illumina reads from *Pgt* Ug99 or *Pgt*21-0 to Ug99 assembly. For **a** and **b** y-axis depicts genome coverage calculated in 1 kb sliding windows and normalized to the mean of coverage of each haplotype. Genome coverage shows a normal distribution for self-mapping. Read cross-mapping also shows a normal distribution for haplotype A of Ug99 and *Pgt*21-0 which indicates high sequence similarity. In contrast, a skewed distribution to low genome coverage occurs in the B and C haplotype comparison due to high sequence divergence. **c** Numbers of homozygous SNPs and MNPs called for various extracted Illumina read sets mapped against the *Pgt Pgt*21-0 (grey) or Ug99 (black) reference genome assemblies. Illumina reads were first mapped at high stringency to the corresponding reference genome and then uniquely mapped reads from each haplotype were extracted and used for variant calling. The low number of SNPs detected in the inter isolate comparisons of haplotype A in contrast to the high number of SNPs identified in the B or C haplotype, supports the close identity of A haplotypes in both isolates.

**Supplementary Fig. 4.**
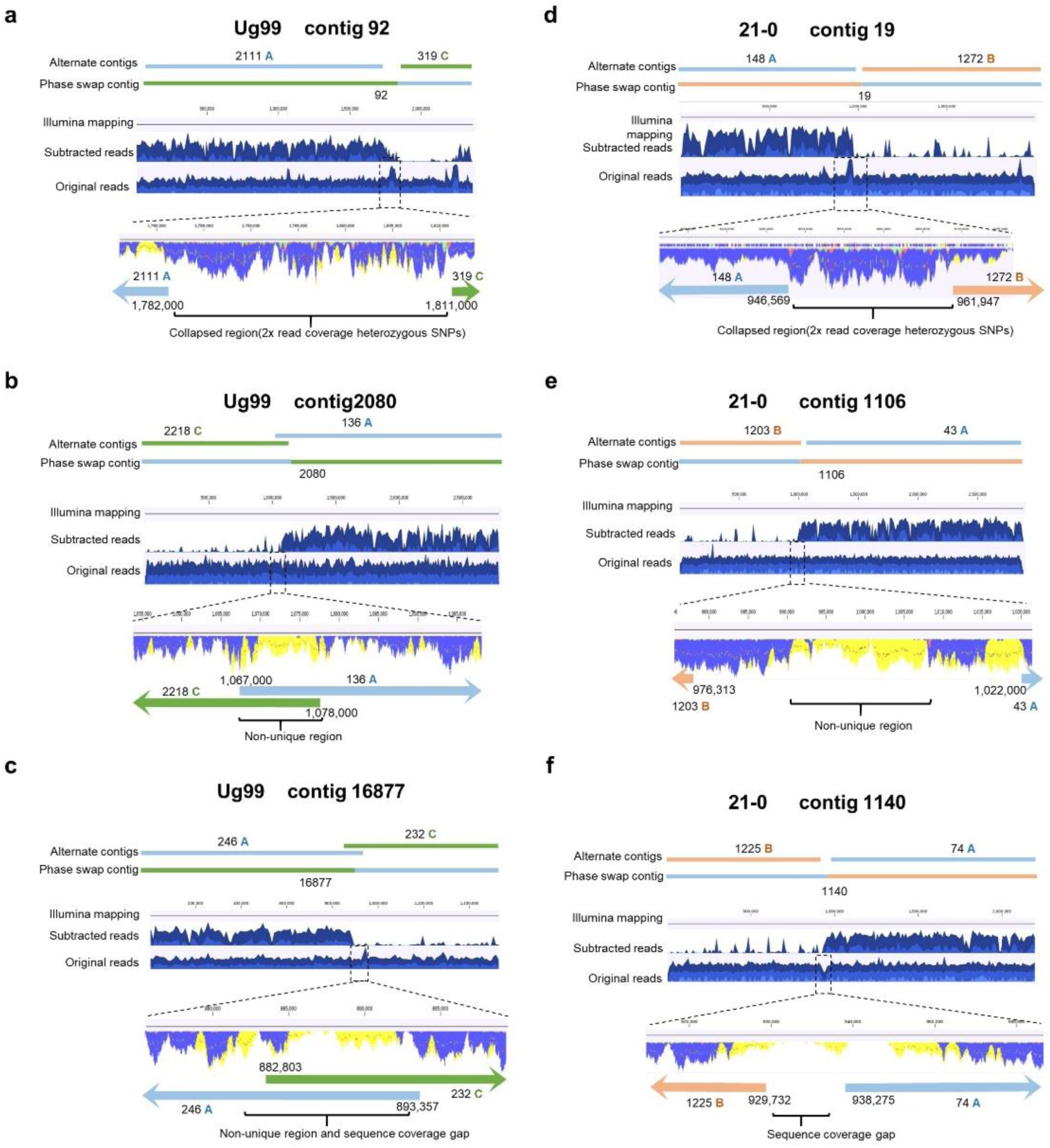
Examples illustrating the detection and manual curation of phase swap contigs in the *Pgt Pgt*21-0 and Ug99 genome assemblies. **a** to **f**, The top of each figure shows chimeric contigs and alternate contigs colour-coded according to haplotype assignment. The next two tracks show read coverage graphs of subtracted reads and original reads across the phase swap contigs visualized in CLC Genomics Workbench browser (see read subtraction procedure Supplementary Figure 4). Zoomed in regions (dotted boxes) show coverage graphs for the phase swap junction regions. Coloured bars indicate SNP frequencies in the underlying reads, and yellow shading indicates non-uniquely mapped reads. Coloured arrows at the bottom shows alignment positions of the alternate contigs to this region with the endpoint coordinates indicated. These examples illustrate scenarios indicative of assembly errors due to collapsed assembly regions showing double overage with heterozygous SNPs (**a, d**), non-uniquely mapped repeats (**b, e**) or coverage gaps after Illumina read mapping (**c, f**).

**Supplementary Fig. 5.**
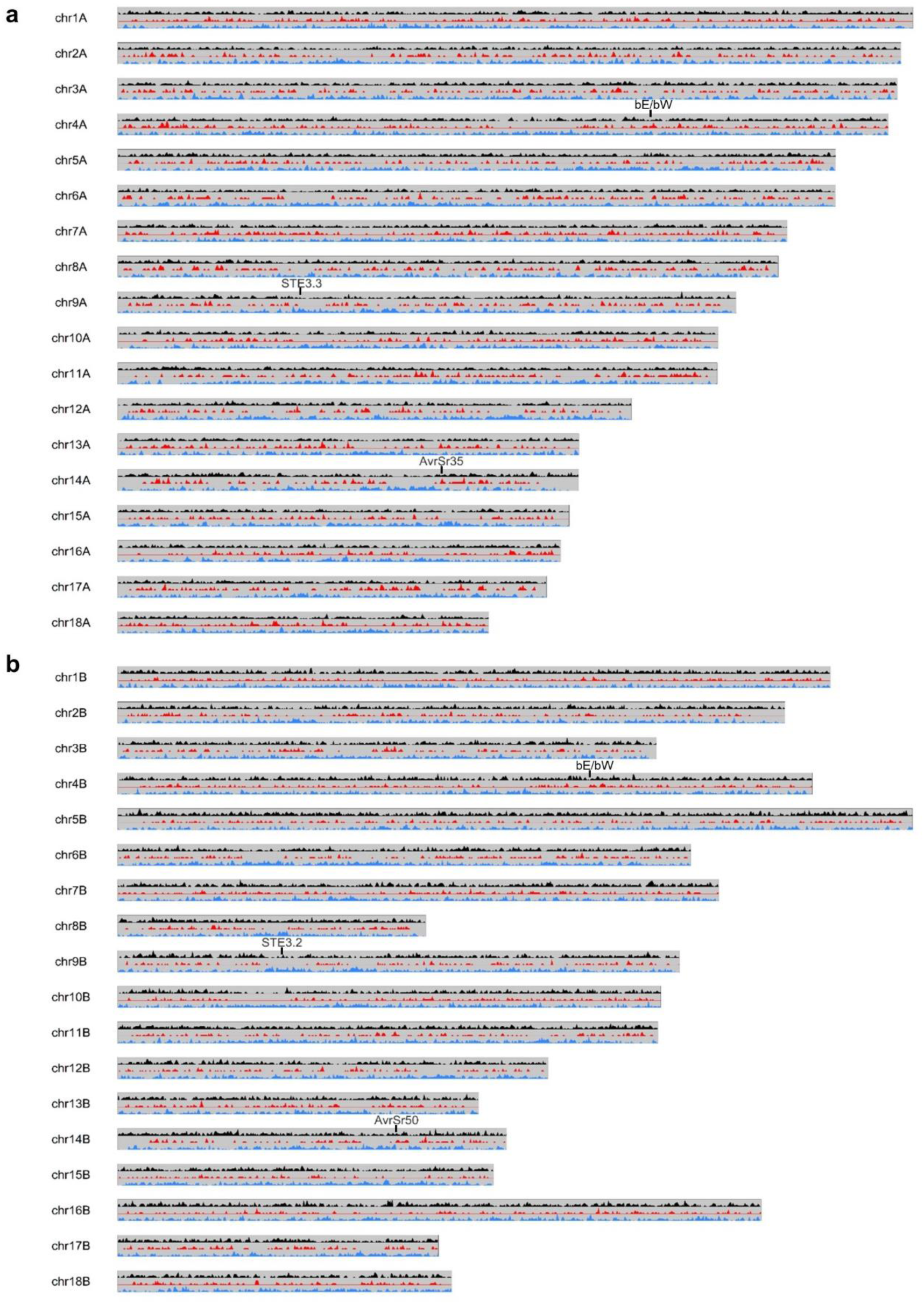
Gene and repeat density plots for homologous chromosomes in haplotypes A and B of *Pgt Pgt*21-0. Top two tracks show density of genes encoding non-secreted (black) or secreted proteins (red) along the chromosomes. Bottom graph shows density of repeat elements (blue). Positions of *bE/bW, STE3.2*, *STE 3.3*, *AvrSr50* and *AvrSr35* genes are indicated.

**Supplementary Fig. 6.**
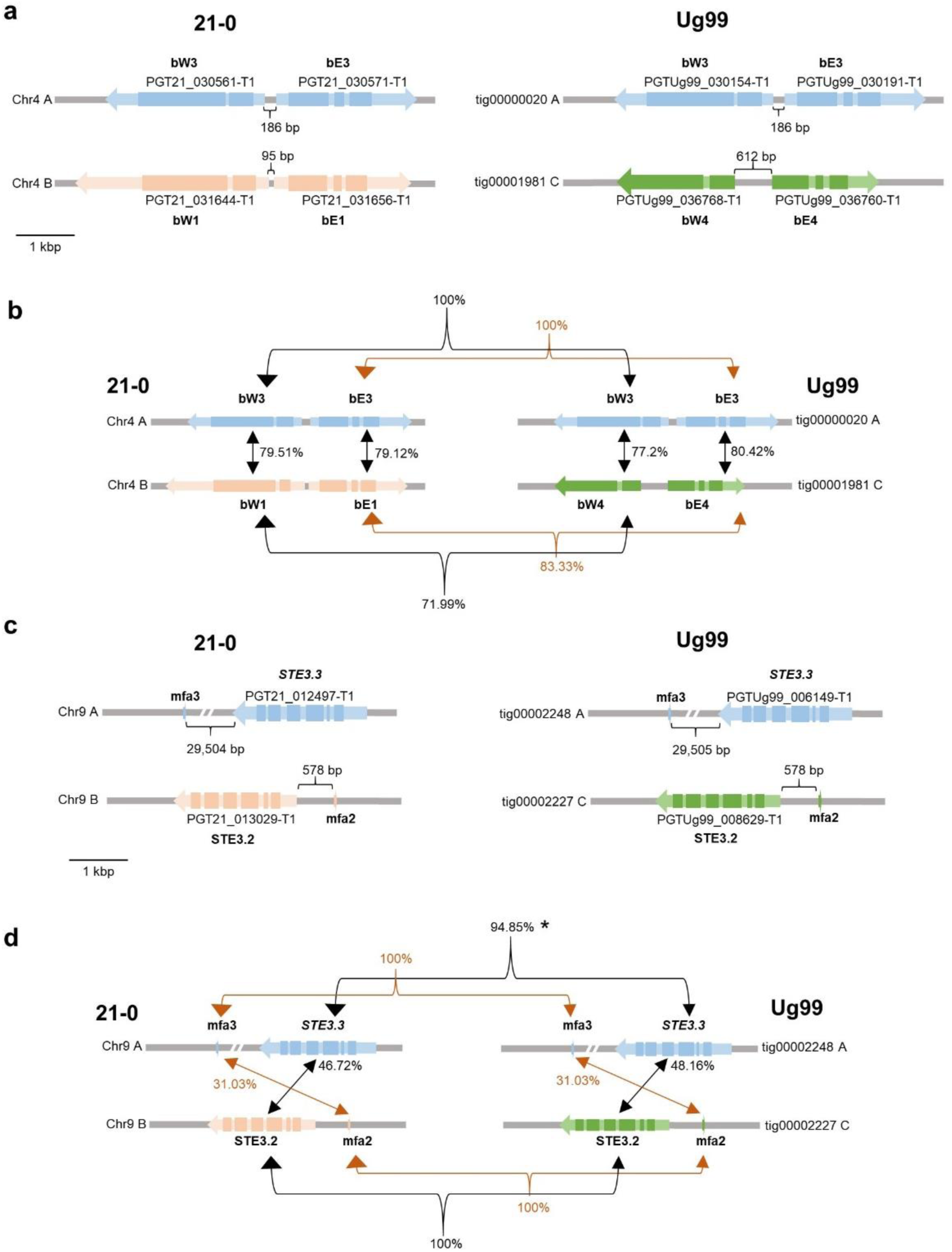
Structure of mating type loci in *Pgt Pgt*21-0 and Ug99. The predicted *a* and *b* loci are on separate chromosomes, consistent with a heterothallic nature controlled by two unlinked loci. **a,** Divergent orientation of the *bE*/*bW* genes from the *b* mating-type locus on chromosome 4. The gene transcripts and orientations are depicted by light coloured arrows and coding sequences by the darker boxes. Colour coding represents the three haplotypes (A=blue, B=orange, C=green). The distances between predicted gene models is shown. The *bW1*/*bE1* allele is identical to the *Pgt bE1*/*bW1* allele previously identified in a North American isolate 75-36-700-3^79^. The bE2/bW2 allele from 75-36-700-3 was not present in either isolate, which instead contained two additional novel alleles, *bE3*/*bW3* and *bE4*/*bW4* alleles, indicating that this locus is multi-allelic in *Pgt*. **b,** Percentage amino acid identity between predicted proteins encoded by *bE* and *bW* alleles within and between *Pgt* isolates. **c,** Arrangement of the pheromone peptide encoding genes (*mfa2* or *mfa3*) and pheromone mating factor receptors (*STE3.2* and *STE3.3*) at the predicted *a* mating type locus. The two alleles of the *a* locus on chromosome 9 contain either the STE3.2 (B and C haplotypes) or STE3.3 (A haplotype) predicted pheromone receptor genes from CRL 75-36-700-3^79^ in both isolates, consistent with a binary recognition system. d, Percentage amino acid identity between pheromone peptide and receptor alleles within and between *Pgt* isolates. The *STE3.3* allele in Ug99 is identical to that in *Pgt*21-0 except for a 1 bp deletion causing a frameshift and replacement of the last 48 amino acids by an unrelated 24 amino acid sequence resulting in reduced amino acid identity (*).

**Supplementary Fig. 7.**
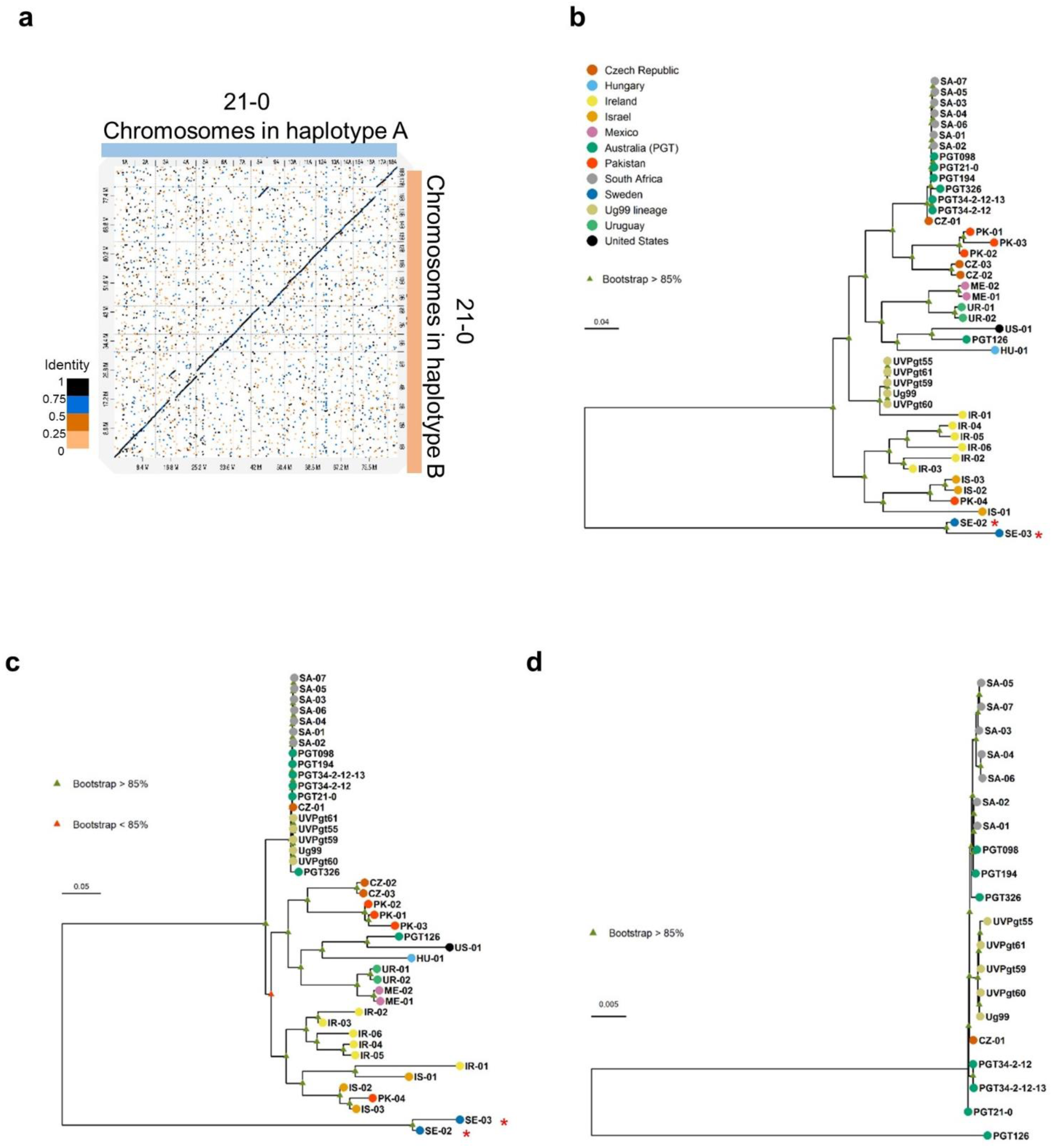
Phylogenetic analysis of Pgt isolates from diverse countries of origin using a RAxML model. **a,** Dot plot of sequence alignment of *Pgt Pgt*21-0 chromosome pseudomolecules of haplotypes A and B. Two translocation events, one between chromosomes 3 and 5 and one between chromosomes 8 and 16, are evident. **b,** Dendrogram inferred using biallelic SNPs detected against the complete diploid genome assembly of *Pgt* Ug99. **c,** Dendrogram inferred from SNPs detected in haplotype A of Ug99. **d,** Dendrogram inferred using biallelic SNPs in haplotype A of *Pgt Pgt*21-0 for the South African, Australian and Ug99 lineage isolates that share the A haploid genome with *Pgt* 126 included as an outgroup. Colour key in panel **b** indicates country of origin for all dendrograms. Scale bar indicates number of nucleotide substitutions per site. Red asterisks indicate *P. graminis* f. sp. *avenae* isolates.

**Supplementary Fig. 8.**
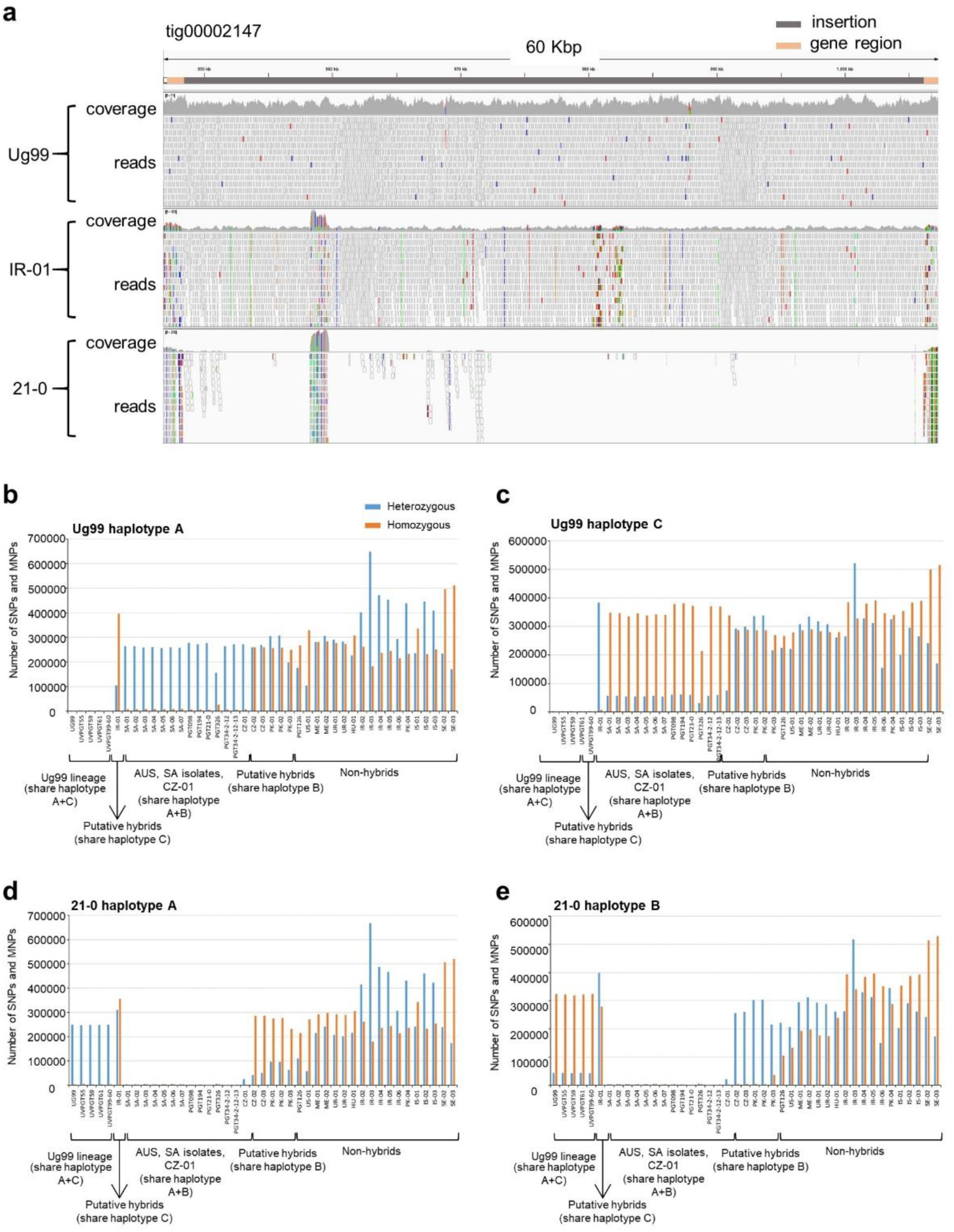
Putative *Pgt* hybrids that share the B or C haplotypes of *Pgt Pgt*21-0 and Ug99, respectively. **a,** Genome browser view in IGV of a 60 kbp genomic region in haplotype C of Ug99. The top bar shows the *AvrSr35* coding sequences (orange) flanking a 57 kbp-insert (grey). Following tracks illustrate coverage and Illumina read alignments of Ug99, IR-01, and *Pgt*21-0. In contrast to *Pgt*21-0, the genome of IR-01 contains a sequence similar to the 57 kbp insert in Ug99. **b** to **f,** Bar graphs show number of homozygous (orange) and heterozygous (blue) SNPs and MNPs called against the *Pgt*21-0 or Ug99 A, B and C haplotypes from Illumina read data for 43 *Pgt* isolates used for phylogenetic analysis. Read mapping patterns to each haplotype vary according to the presence or absence of either the A, B or C haplotypes in each isolate. Considering read mapping to the *Pgt*21-0 reference first; for *Pgt*21-0 and the other clonal Australian and South African isolates containing both A and B haplotypes, reads from the A nucleus will map to the A genome and reads from the B nucleus will map to the B genome. A very low number of homozygous SNPs are therefore detected that represent accumulated mutations as this lineage has evolved. For Ug99 and other A+C haplotype isolates in this clonal group, reads from the A nucleus will map to the A genome and again any new mutations appear as a small number of homozygous SNPs in the A genome data set. However, reads from the C genome can map to either the A or B genomes according to sequence similarity. If they map to the B genome they will give rise to homozygous SNPs representing divergence between the B and C genomes. If they map to the A genome they give rise to heterozygous SNPs because A genome reads are already mapped to these regions. Thus, we see a small number of homozygous SNPs on the A genome, and a large number of homozygous SNPs on the B genome and a similarly large number of heterozygous SNPs on the A genome. Other isolates that are not hybrids derived from race 21 will have two nuclei that are neither A nor B, and reads from these can map to either the A or B genomes giving rise to high numbers of both heterozygous and homozygous SNPs on each haplotype. Thus, the variation in these patterns of heterozygous and homozygous SNPs on the different haplotypes are indicative of different hybrid relationships. The patterns for CZ-02,03 and PK-01,02 are consistent with them containing haplotype B, with many heterozygous but very few homozygous SNPs called on this haplotype, while IR-01 shows a similar pattern on haplotype C, again indicating that it contains a very similar haplotype.

**Supplementary Table 2.**
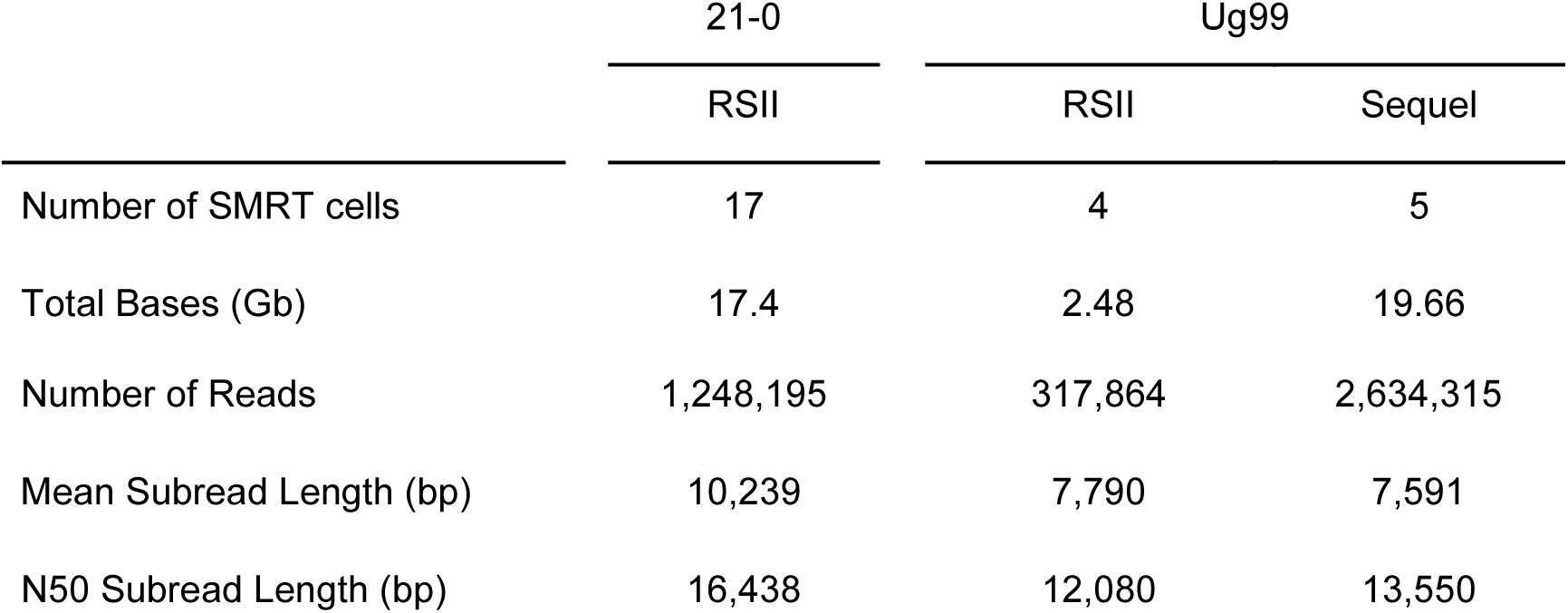
Summary statistics for SMRT sequencing and raw read metrics.

|  | 21-0 | Ug99 |  |
| --- | --- | --- | --- |
|  | RSII | RSII | Sequel |
| Number of SMRT cells | 17 | 4 | 5 |
| Total Bases (Gb) | 17.4 | 2.48 | 19.66 |
| Number of Reads | 1,248,195 | 317,864 | 2,634,315 |
| Mean Subread Length (bp) | 10,239 | 7,790 | 7,591 |
| N50 Subread Length (bp) | 16,438 | 12,080 | 13,550 |

**Supplementary Table 3.** Summary statistics of Illumina sequencing of *Pgt* isolates in the Ug99 lineage.

| Isolate (Pathotype) | 150 bp Paired-End Reads | Yield (Mbp) | Mean Quality Score |
| --- | --- | --- | --- |
| UVPgt55 (TTKSF) | 40,048,658 | 12,094 | 31.53 |
| UVPgt59 (TTKSP) | 27,204,289 | 8,216 | 31.14 |
| UVPgt60 (PTKST) | 36,665,018 | 11,073 | 31.56 |
| UVPgt61 (TTKSF)* | 36,605,359 | 11,055 | 31.57 |
| Ug99 (TTKSK) | 35,674,381 | 10,773 | 31.48 |

\* Virulent on resistance gene *Sr9h*

**Supplementary Table 4.** Assembly metrics and quality analysis

| Parameters | 21-0 | Ug99 |
| --- | --- | --- |
| No. of contigs | 410 | 514 |
| No. of contigs ≥ 50,000 bp | 249 | 333 |
| Total length (Mbp) | 176.9 | 176.0 |
| Total length ≥50000bp (Mbp) | 171.8 | 170.4 |
| Size of Largest contig (Mbp) | 5.96 | 4.40 |
| N50 (Mbp) | 1.26 | 0.97 |
| GC (%) | 43.5 | 43.5 |
| No. of contigs with telomeres | 69 | 26 |
| % of complete BUSCOs | 95.8 | 95.6 |
| % single-copy BUSCOs | 8.3 | 8.9 |
| % duplicated BUSCOs | 87.5 | 86.7 |
| % of fragmented BUSCOs | 1.9 | 1.9 |
| % of missing BUSCOs | 2.3 | 2.5 |
| No. of Bins | 44 | 62 |
| Total length in bins (Mbp) | 169 | 165 |
| No. contigs in bins | 225 | 276 |

**Supplementary Table 6.** Intra and inter-isolate sequence comparison of the *AvrSr50* chromosome haplotypes in *Pgt* Ug99 and *Pgt*21-0.

| Isolate comparison | Sequence similarity |  |  |  |  |  |  | Structural variation (SV) |  |  |
| --- | --- | --- | --- | --- | --- | --- | --- | --- | --- | --- |
|  | Bases aligned (%) | Average identity of alignment blocks (%) | Overall identity (%) | Divergence of aligned blocks (%) | Total SNPs | SNPs/kbp | Indels | Number of variants >50bp | Total size of variants |  |
|  |  |  |  |  |  |  |  |  | Mbp | % of chromosome |
| Ug99 A vs *Pgt*21-0 A | 99.8 | 99.93 | 99.73 | 0.07 | 307 | 0.10 | 820 | 29 | 0.17 | 2.56 |
| Ug99 A vs Ug99 C | 70.82 | 95.12 | 67.36 | 4.88 | 52,839 | 25.28 | 34,563 | 167 | 1.3 | 22.03 |
| 21-0 A vs *Pgt*21-0 B | 73.12 | 95.50 | 69.83 | 4.50 | 57,463 | 22.03 | 37,070 | 190 | 1 | 14.03 |
| Ug99 C vs *Pgt*21-0 B | 78.36 | 97.27 | 76.22 | 2.73 | 33,655 | 14.56 | 21,766 | 137 | 1.09 | 16.74 |
| Ug99 A vs *Pgt*21-0 B | 71.87 | 94.98 | 68.26 | 5.02 | 55,310 | 26.07 | 35,314 | 187 | 1.25 | 19.2 |
| 21-0 A vs Ug99 C | 64.26 | 95.11 | 61.12 | 4.89 | 54,593 | 23.82 | 35,340 | 150 | 0.97 | 14.8 |

<sup>a</sup> First listed isolate served as reference and second listed isolate served as query for the analysis.
<sup>b</sup> Overall identity is average identity of alignment block multiplied by the proportion of bases aligned.

**Supplementary Table 9.**
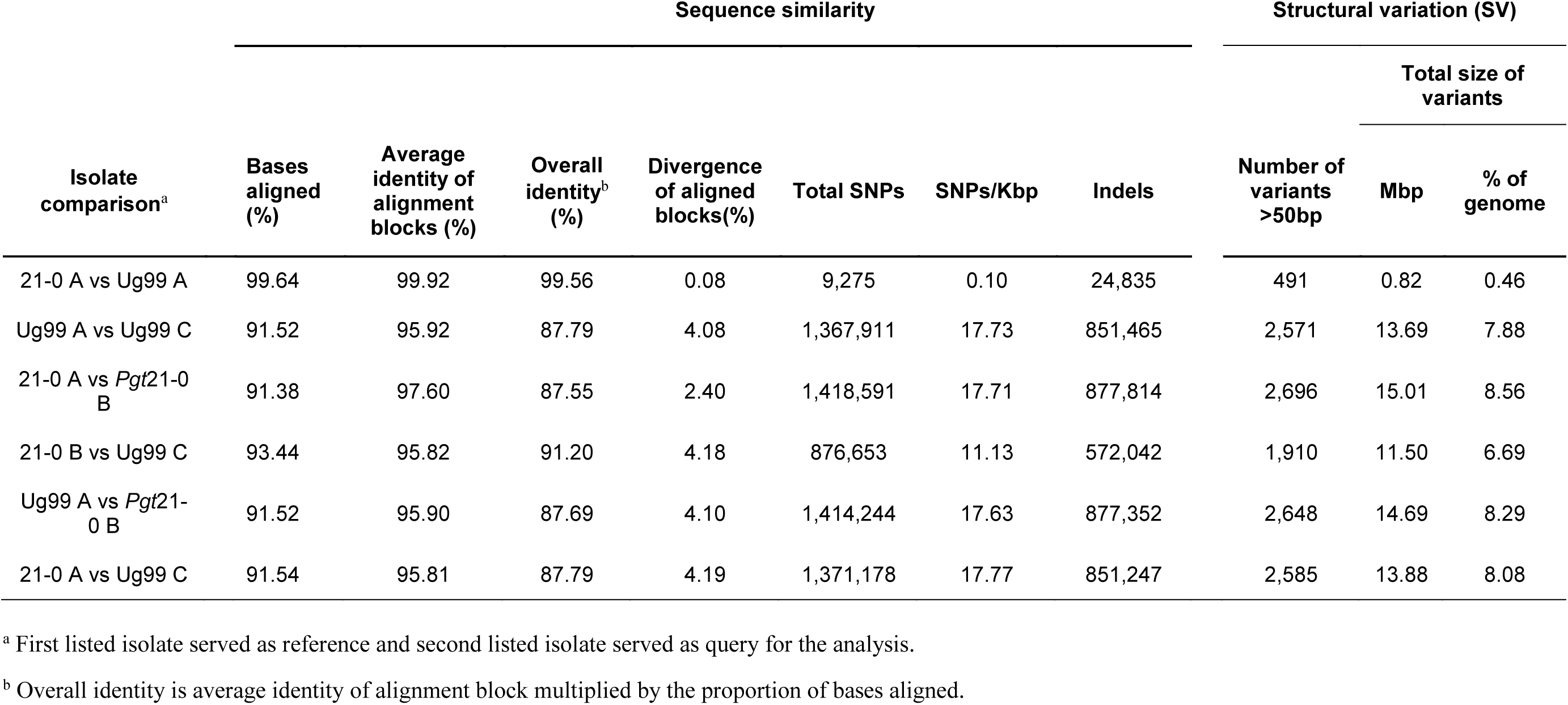
Intra- and inter-isolate sequence comparison of entire haplotypes in *Pgt* Ug99 and *Pgt*21-0.

| Isolate comparison<sup>a</sup> | Sequence similarity |  |  |  |  |  |  | Structural variation (SV) |  |  |
| --- | --- | --- | --- | --- | --- | --- | --- | --- | --- | --- |
|  | Bases aligned (%) | Average identity of alignment blocks (%) | Overall identity<sup>b</sup> (%) | Divergence of aligned blocks(%) | Total SNPs | SNPs/Kbp | Indels | Number of variants >50bp | Total size of variants |  |
|  |  |  |  |  |  |  |  |  | Mbp | % of genome |
| 21-0 A vs Ug99 A | 99.64 | 99.92 | 99.56 | 0.08 | 9,275 | 0.10 | 24,835 | 491 | 0.82 | 0.46 |
| Ug99 A vs Ug99 C | 91.52 | 95.92 | 87.79 | 4.08 | 1,367,911 | 17.73 | 851,465 | 2,571 | 13.69 | 7.88 |
| 21-0 A vs *Pgt*21-0 B | 91.38 | 97.60 | 87.55 | 2.40 | 1,418,591 | 17.71 | 877,814 | 2,696 | 15.01 | 8.56 |
| 21-0 B vs Ug99 C | 93.44 | 95.82 | 91.20 | 4.18 | 876,653 | 11.13 | 572,042 | 1,910 | 11.50 | 6.69 |
| Ug99 A vs *Pgt*21-0 B | 91.52 | 95.90 | 87.69 | 4.10 | 1,414,244 | 17.63 | 877,352 | 2,648 | 14.69 | 8.29 |
| 21-0 A vs Ug99 C | 91.54 | 95.81 | 87.79 | 4.19 | 1,371,178 | 17.77 | 851,247 | 2,585 | 13.88 | 8.08 |

<sup>a</sup> First listed isolate served as reference and second listed isolate served as query for the analysis.
<sup>b</sup> Overall identity is average identity of alignment block multiplied by the proportion of bases aligned.

**Supplementary Table 11.**
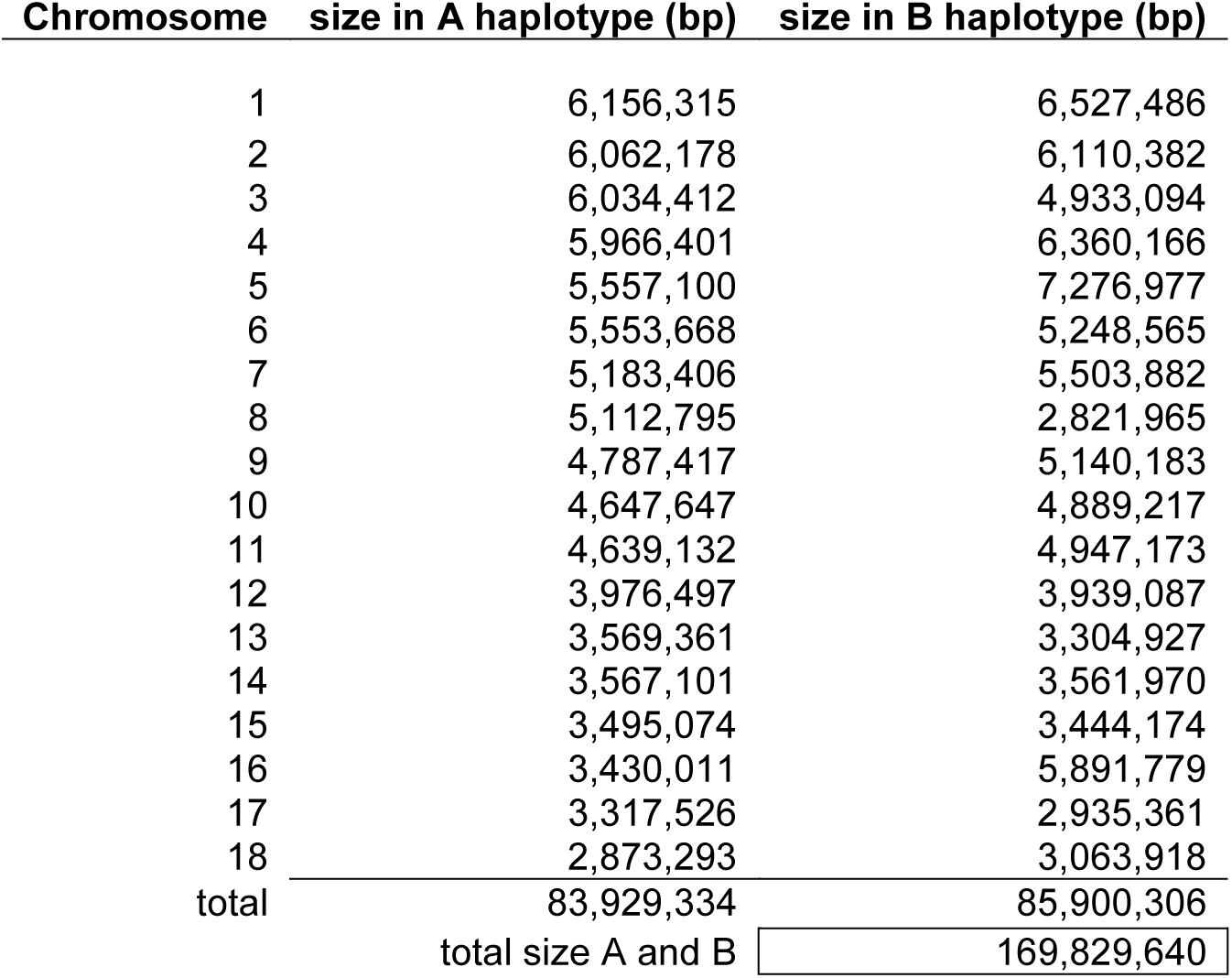
Assignment of contigs to chromosomes in *Pgt*21-0

| **Chromosome** | **size in A haplotype (bp)** | **size in B haplotype (bp)** |
| --- | --- | --- |
| 1 | 6,156,315 | 6,527,486 |
| 2 | 6,062,178 | 6,110,382 |
| 3 | 6,034,412 | 4,933,094 |
| 4 | 5,966,401 | 6,360,166 |
| 5 | 5,557,100 | 7,276,977 |
| 6 | 5,553,668 | 5,248,565 |
| 7 | 5,183,406 | 5,503,882 |
| 8 | 5,112,795 | 2,821,965 |
| 9 | 4,787,417 | 5,140,183 |
| 10 | 4,647,647 | 4,889,217 |
| 11 | 4,639,132 | 4,947,173 |
| 12 | 3,976,497 | 3,939,087 |
| 13 | 3,569,361 | 3,304,927 |
| 14 | 3,567,101 | 3,561,970 |
| 15 | 3,495,074 | 3,444,174 |
| 16 | 3,430,011 | 5,891,779 |
| 17 | 3,317,526 | 2,935,361 |
| 18 | 2,873,293 | 3,063,918 |
| total | 83,929,334 | 85,900,306 |
| total size A and B |  | 169,829,640 |

**Supplementary Table 13.** Summary of gene annotation

|  | **21-0** | **Ug99** |
| --- | --- | --- |
| No. of genes including tRNAs | 37,061 | 37,394 |
| No. of protein coding genes | 36,319 | 36,659 |
| - haplotype A | 18,225 (17,786)* | 18,593 |
| - haplotype B or C | 17,919 (17,718)* | 17,621 |
| % of genome covered by genes | 34.9 | 33.6 |
| Mean gene length (bp) | 1,667 | 1,586 |
| No. of secreted protein genes | 6,180 | 6,120 |
| - haplotype A | 3,099 (3,071)* | 3,212 |
| - haplotype B or C | 3,063 (3,046)* | 2,857 |

\*No. of predicted genes in contigs assigned to chromosomes

**Supplementary Table 14.** Shared and unique gene content between *Pgt* haplotypes

|  | Haplotypes compared |  |  | Haplotypes compared |  |  |
| --- | --- | --- | --- | --- | --- | --- |
|  | 21-0 A | 21-0 B | Ug99 C | Ug99 A | 21-0 B | Ug99 C |
| Unique genes | 3,369<br>(18%) | 2,668<br>(15%) | 2,950<br>(17%) | 3,529<br>(19%) | 2,774<br>(15%) | 2,950<br>(17%) |
| genes shared with one<br>other haplotype | 2,492<br>(14%) | 3,165<br>(18%) | 2,543<br>(14%) | 2,976<br>(16%) | 2,664<br>(15%) | 2,827<br>(15%) |
| genes shared in three<br>haplotypes | 12,364<br>(68%) | 12,086<br>(67%) | 12,128<br>(69%) | 12,088<br>(65%) | 12,481<br>(70%) | 12,082<br>(69%) |
| total genes | 18,225 | 17,919 | 17,621 | 18,593 | 17,919 | 17,621 |

**Supplementary Table 15.**
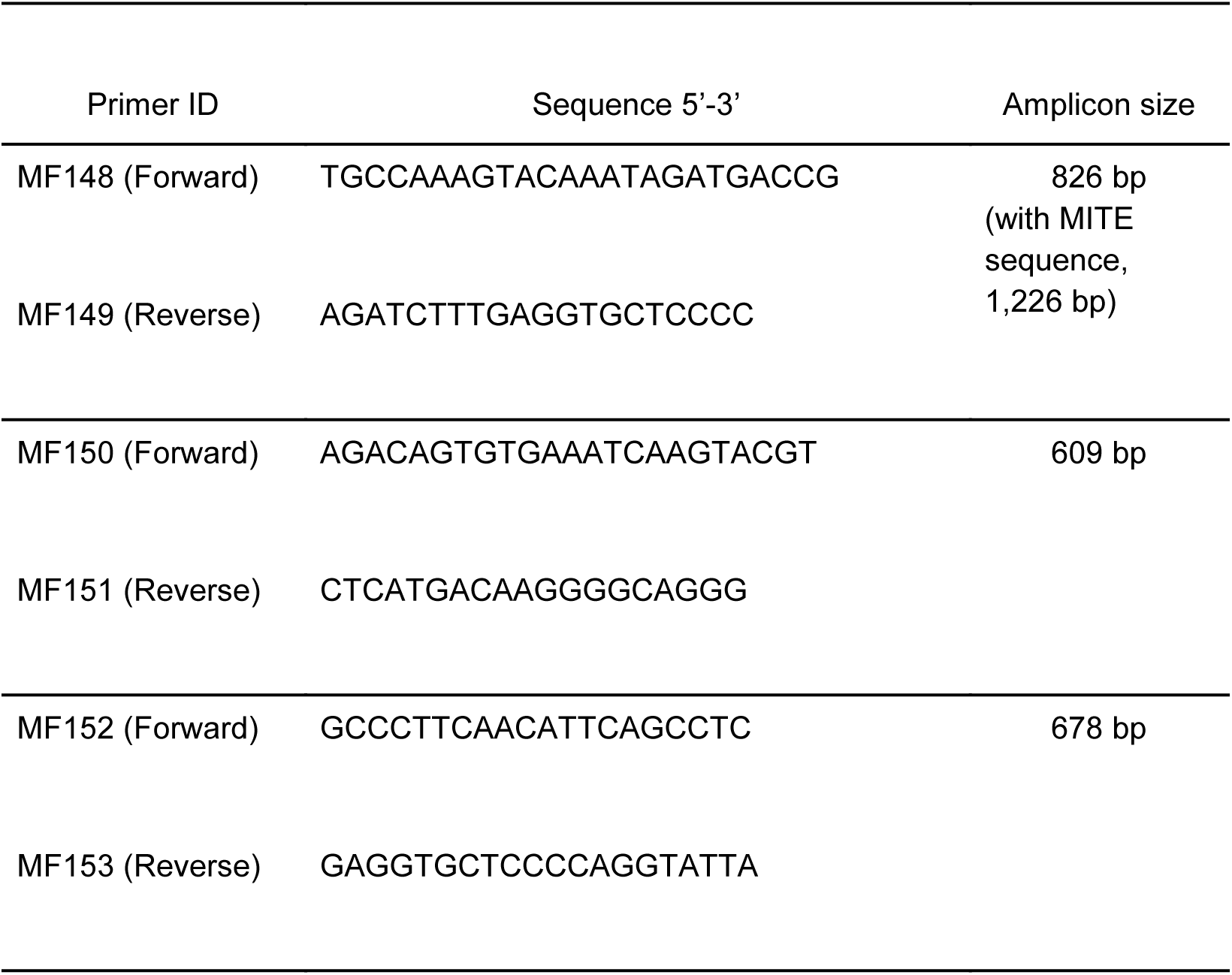
List of primer sequences to amplify flanking and internal regions of the 57 kbp insert in *AvrSr35*.

## Additional files

**Supplementary Table 1.**

Virulence reactions and pathotype assignments of *Pgt* isolates in the Ug99 lineage. Scores are reported based on the North American wheat differential set (excel file).

**Supplementary Table 5.**

Gene synteny output (excel file)

**Supplementary Table 7.**

Summary of karyon assignment before breaking chimeric contigs in *Pgt* Ug99 and *Pgt*21-0 (excel file)

**Supplementary Table 8.**

List of chimeric contigs and breakpoints (separate file)

**Supplementary Table 10.**

Physical linkage of phase swap contigs in the *Pgt Pgt*21-0 assembly to contigs of the same or alternate haplotype within bin or chromosome calculated from Hi-C data (excel file)

**Supplementary Table 12.**

Contigs assigned to chromosomes (excel file)

**Supplementary Table 16.**

Metadata for RNAseq libraries of *Pgt Pgt*21-0 used for training gene models in the annotation pipeline (xcel file).

**Supplementary Table 17.**

Metadata genome coverages after mapping Illumina reads to *Pgt Pgt*21-0 and Ug99 references (excel file)

